# Defining the OrrA regulon and its role in development and antibiotic production in *Streptomyces venezuelae* NRRL B-65442

**DOI:** 10.64898/2026.01.30.702861

**Authors:** Kelly-Rose Tulley, Lucas Balis, Neil A. Holmes, Ainsley D. M. Beaton, Courtney Andrews, Gerhard Saalbach, Rhea Stringer, Govind Chandra, Barrie Wilkinson, Matthew I. Hutchings

## Abstract

2.

*Streptomyces* bacteria have complex life cycles involving hyphal growth, sporulation, and the production of diverse specialised metabolites, including antibiotics. In this study, we investigated the role of the highly conserved orphan response regulator OrrA in *Streptomyces venezuelae* NRRL B-65442. We show that *S. venezuelae* Δ*orrA* mutants are defective in sporulation and, using ChIP-seq, identify five OrrA binding sites *in vivo*. Tandem-mass-tag proteomics revealed that OrrA directly activates two of these putative target genes, *wblA* and *vnz_04640*, a finding consistent with previous work on OrrA in the distantly related *S. coelicolor*. We also demonstrate that deleting *wblA* blocks sporulation and that overexpressing *wblA* restores sporulation in the Δ*orrA* mutant. Additionally, chloramphenicol biosynthesis is upregulated in both the Δ*orrA* and Δ*wblA* mutants compared with the wild type. Taken together, these results indicate that the primary function of OrrA is to regulate WblA production, and that reduced intracellular WblA levels underlie the phenotypes observed in the Δ*orrA* mutant.

**Data summary:** The authors confirm all supporting data, code and protocols have been provided within the article, through supplementary data files and via public databases.

## 4. Introduction

The genus *Streptomyces* comprises more than 1100 verified species and is the type genus for the ancient bacterial phylum Actinomycetota (1). *Streptomyces* species are studied for their complex development and their prolific production of specialised metabolites, which form the basis of ∼55% of clinically used antibiotics (2). The *Streptomyces* life cycle begins with spore germination and outgrowth into a branching, vegetative substrate mycelium which, upon nutritional stress, gives rise to reproductive aerial hyphae that undergo rapid DNA replication and cell division to form chains of unigenomic spores (3). Prior to genome sequencing, classical genetics identified many genes that are involved in development, and these were classified as either *bld* (bald) genes, that are required for aerial hyphae formation, or *whi* (white) genes that are required for sporulation (1). The *bld* mutants form colonies that do not have ‘hairy’ aerial mycelium and thus appear bald when growing on solid agar while the *whi* mutants form white colonies that lack the WhiE spore pigment, either because the aerial hyphae do not undergo cell division to form spores or because the spores do not mature (3).

Two-component systems (TCS) as well as orphan response regulators (RRs) and sensor kinases (SKs) have been heavily implicated in the regulation of the *Streptomyces* life cycle and their specialised metabolism (4,5). This includes the RR MtrA which is a master regulator of specialised metabolism in *Streptomyces* species (6–9). MtrA is part of a conserved three-component system that includes its cognate SK MtrB and an accessory lipoprotein LpqB (10). Recent work reported that MtrAB senses osmotic stress in *S. venezuelae* NRRL B-65442 and activates production of the compatible solute ectoine and triggers entry into sporulation (11). An orphan RR designated OrrA (12) is encoded within the same region and the *orrA* gene (*sco3008*) is separated from the *mtrAB-lpqB* operon (*sco3013-11*). Deletion of *orrA* in *Streptomyces coelicolor* M145 resulted in a bald mutant that cannot form aerial hyphae or spores, even after prolonged incubation (13). Only two OrrA target genes were identified in *S. coelicolor* using ChIP-seq, namely *sco1375*, which encodes a small protein of unknown function, and *wblA* which encodes a member of the WhiB-like family of 4Fe-4S cluster containing transcription factors (14–16). RNA-seq analysis showed that OrrA is required to activate the expression of *sco1375* and *wblA* and the *S. coelicolor ΔwblA* and *ΔorrA* mutants share very similar phenotypes. However, over-expression of either gene did not fully complement the *S. coelicolor* Δ*orrA* mutant.

In this work we show that OrrA activates the production of WblA and the SCO1375 homologue Vnz_04640 in the model organism *S. venezuelae* NRRL B-65442, which is distantly related to *S. coelicolor. S. venezuelae* NRRL B-65442 forms green spores and has emerged as a model organism because of its genetic tractability, rapid growth and the ability to complete its life cycle to sporulation in liquid growth medium (17,18). Three additional putative OrrA target promoters were identified here using ChIP-seq and we used a DNA footprinting technique called ReDCaT SPR to identify OrrA binding sites at four of these five promoters. However, of the five putative OrrA target genes identified, proteomics analysis showed that only two of the gene products were significantly affected in abundance by loss of OrrA, namely WblA and Vnz_04640. *S. venezuelae ΔorrA* and *ΔwblA* mutants are both defective in sporulation and over produce the antibiotic chloramphenicol suggesting the phenotypes observed in the *ΔorrA* strain are the result of a reduction in WblA abundance. Consistent with this, over-expression of *wblA* in the *ΔorrA* mutant restored sporulation. We conclude that OrrA targets and function are conserved between *S. coelicolor* M145 and *S. venezuelae* NRRL B-65442, namely to control the expression of *wblA*, which encodes a key regulator of development and a global repressor of antibiotic biosynthesis (19).

## 5. Methods

### Bacterial strains

Wild-type *Streptomyces venezuelae* NRRL B-65542 and isogenic mutant strains were grown in MYM (Maltose + Yeast Extract + Malt Extract) liquid medium and on MYM agar. *Escherichia coli* strains were grown in LB (Lennox Broth) and on LB agar.

Media recipes plus written and video protocols for culturing *Streptomyces* bacteria, making spore preparation and inter-species conjugation can be found at http://actinobase.org and in the *Streptomyces* Practical Genetics manual which is free to download here: https://streptomyces.org.uk/PracticalStreptomycesGenetics.pdf (20). All the bacterial strains, plasmids and primers used in this study are listed in Tables S1-S3.

### CRISPR/Cas9-mediated genome editing

The *orrA* gene was deleted using pCRISPomyces-2 following the published protocol (21) using gene-specific guide RNAs and repair templates (see Table S3 for primers). A detailed protocol is also available at http://actinobase.org (22).

### Chloramphenicol Analysis

Chloramphenicol was measured according to Som et al., 2017 (6). Strains were grown in biological triplicates in 30ml MYM liquid medium for 24 hours. An aliquot of 750 μL was taken from each culture and 750 μL of ethyl acetate was added. After shaking for 10 min, the extracts were centrifuged, and the organic phase (500 μL) was transferred to a 1.5 mL glass vial. The solution was concentrated under reduced pressure and the residual extract dissolved in 500 μL of a mixture of methanol:water (2:3). For quantification 40 μL of each sample was subjected to analytical HPLC. Analysis was performed on an Agilent 1290 Infinity II LC System (Agilent Technologies). A Gemini^®^ 3.0 μm NX-C18 150 x 4.6 mm column was used. Chloramphenicol was eluted using mobile phase A (water containing 0.1% formic acid) and mobile phase B (100% methanol) using the following multistep gradient at a flow rate 800 μL min^-1^: 0 min, 10%B; 14 min, 100%B; 18 min, 100%B; 18.5 min, 10%B; 19.5 min, 10%B. Stock solutions of chloramphenicol (Sigma Aldrich) for calibration were prepared between a range of 0.01 μg mL^-1^ and 0.1 mg mL^-1^.

Generation of the calibration curve and quantification of chloramphenicol was accomplished by integration of the chloramphenicol signal at 273 nm.

### Chromatin Immunoprecipitation Sequencing (ChIP-seq)

*S. venezuelae* wild-type and Δ*orrA* complemented with pSS170 + *orrA*-N-FLAG (Tables S1 and S2) were inoculated from single colonies onto sterile cellophane disks placed on MYM agar and incubated at 30°C for 18 and 48 hours. Under sterile conditions, lawns were scraped into 50 mL tubes, and cross-linking was performed by adding 10 mL of 1% (v/v) formaldehyde in PBS for 30 minutes at 30°C with gentle shaking (90 rpm). Cross-linking was quenched by replacing the formaldehyde with 10 mL of 0.5 M glycine and incubating for 5 minutes at room temperature. Following glycine removal, the biomass was washed twice with 25 mL of ice-cold PBS (pH 7.4) and flash frozen at –80 °C. Unless otherwise stated, all buffers used during extraction contained EDTA-free protease inhibitors (2 tablets per 10 mL). Cell lysis was performed by resuspending pellets in 2 mL of lysis buffer (10 mM Tris-HCl pH 8.0, 50 mM NaCl, 10 mg/mL lysozyme) and incubating at 37 °C for 30 minutes. DNA fragmentation was initiated by adding 1 mL of immunoprecipitation (IP) buffer (100 mM Tris-HCl pH 8.0, 500 mM NaCl, 1% Triton X-100) and mixing by pipetting. Samples were sonicated on ice (20 cycles at 50 Hz, 10 second pulse with 1-minute rest between pulses). Fragmentation efficiency was checked by phenol–chloroform extraction of 25 μL crude lysate mixed with 75 μL TE buffer (10 mM Tris-HCl pH 8.0, 1 mM EDTA), followed by RNase A treatment (2 μL of 1 mg/mL) and agarose gel electrophoresis. Ideal fragment sizes ranged between 200–1000 bp. The remaining lysate was cleared by centrifugation at 1968xg for 15 minutes at 4 °C. Anti-FLAG M2 agarose beads (60 μL per sample) were pre-washed twice in 2.5 mL of 0.5× IP buffer and 60 μL was incubated with the cleared lysate overnight at 4°C on a vertical rotor. Beads were subsequently washed four times with 0.5× IP buffer (10 minutes per wash at 4 °C). Bound DNA was eluted by adding 100 μL elution buffer (50 mM Tris-HCl pH 8.0, 10 mM EDTA, 1% SDS) and incubating at 65 °C overnight. A second elution step with 50 μL elution buffer was performed for 5 minutes at 65 °C. Combined eluates were treated with 2 μL proteinase K (20 mg/mL) at 55 °C for 2 hours, followed by phenol–chloroform extraction and purification using a QIAquick column (Qiagen). DNA was eluted in 50 μL EB buffer (10 mM Tris-HCl, pH 8.5); 5 μL was retained for quantification using Nanodrop and Qubit, and the remainder stored for sequencing. Samples were sent to Genewiz® for paired end 150bp Illumina sequencing, generating ∼6 Gb of raw data at ∼20 million read pairs per sample. Sequencing data were processed as described previously (23).

### Reuseable DNA Capture followed by Surface Plasmon Resonance (ReDCaT SPR)

The double stranded oligonucleotide probes used for the *in vitro* DNA binding studies are shown in Table S4. ReDCaT SPR was performed as described previously (24) using the purified DNA binding domain of *S. venezuelae* OrrA.

### Sample Preparation for Tandem Mass Tag (TMT) proteomics

*S. venezuelae* strains were streaked from single colonies onto MYM (pH 7.3) agar plates on top of sterile cellophane disks and incubated at 30 °C to the desired time points. The cellophane disks were then removed, and the mycelium was scraped into 50 mL tubes containing 10 mL cell lysis buffer (2 % SDS, 50 mM TEAB Buffer pH 8.0, 150 mM NaCl and EDTA-free protease inhibitor tablets, supplied by Roche). Samples were vortex vigorously to disrupt the mycelial mass and minimize clumping, then boiled for 10 minutes. Following boiling, samples were then sonicated using four 20 second pulses at 10 microns, with 1-minute intervals between each cycle. Debris was removed by centrifugation at 4,000 x g for 30 min at room temperature.

The clarified supernatant was transferred to a fresh tube, and protein concentration measured using the Qubit Protein Assay. The volume containing 1 mg protein was aliquoted to a 15 mL Falcon tube. Proteins were precipitated with chloroform/methanol (25). Samples were processed by the John Innes Centre proteomics platform, as described previously (26).

Detailed proteomic methods are provided in the supplementary information.

### Cryo-Scanning Electron Microscopy

Conducted by the Bioimaging facility at the John Innes Centre by Rhea Stringer. Samples were mounted on an aluminium stub using Tissue-TekR (BDH Laboratory Supplies, Poole, England). The stub was immediately plunged into liquid nitrogen slush (approximately −210 °C) to cryo-preserve the material. The frozen sample was then transferred to the cryo-stage of an ALTO 2500 cryo-transfer system (Gatan, Oxford, England) attached to an FEI Nova NanoSEM 450 (FEI, Eindhoven, The Netherlands). Surface frost was sublimated at −95 °C for 4 minutes, after which the sample was sputter-coated with platinum for 120 seconds at 10 mA, at a temperature below −110 °C. The sample was then transferred to the main chamber of the microscope, where the cryo-stage was maintained at approximately −125 °C. Imaging was performed at 3 kV, and digital TIFF files were recorded.

### Phylogenetics

To generate the phylogenetic tree, all Actinomycetota reference genomes with a completeness level of “chromosome” or better were downloaded from the NCBI database and the RpoB and OrrA protein sequences were identified using reciprocal BLAST searching with the *Streptomyces venezuelae* NRRL B-65442 (GCF_001886595.1) gene used as the query. All resulting RpoB sequences were aligned using MAFFT (mafft --auto input.faa > output.fasta) (27), the resulting alignment was imported into the Galaxy EU server (28) and FastTree was used with default settings plus LG+CAT model selected to generate a maximum likelihood phylogenetic tree. Subsequently a presence/absence of OrrA, WblA and AdpA was annotated using iTol (29).

## 6. Results and Discussion

### *S. venezuelae ΔorrA* is defective in sporulation

Cas9-mediated genome editing was used to make an in-frame deletion of the *orrA* gene in *S. venezuelae* NRRL B-65442. After two days growth on MYM agar the wild-type strain forms its characteristic, green-pigmented spores while the *ΔorrA* mutant does not sporulate (Figure 1), which is consistent with the phenotype reported for *S. coelicolor* Δ*orrA* (13). This phenotype is maintained if the colonies are restreaked within three days of growth on MYM agar or used within three days to inoculate 2xYT broth to grow liquid cultures for glycerol freezer stocks. However, prolonged growth (≥5 days) results in heterogeneous colonies, some of which form aerial mycelium and spores (Figure S1).

**Figure 1.**
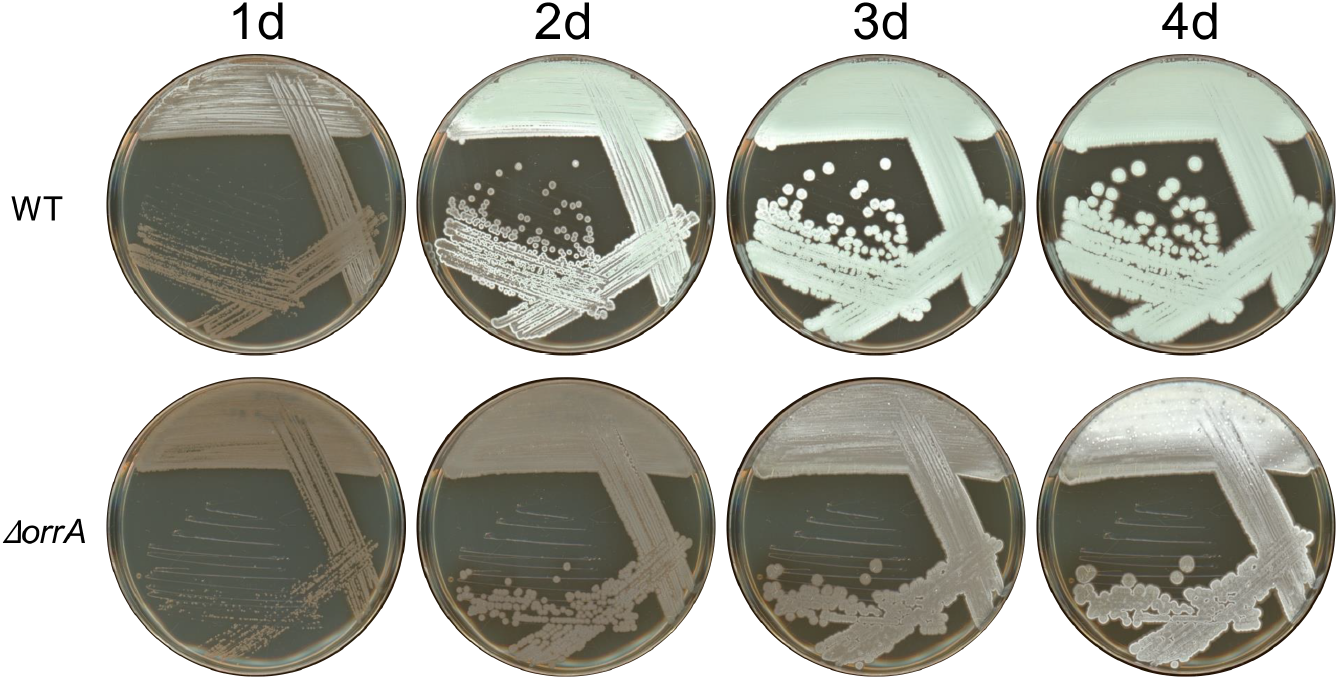
*S. venezuelae ΔorrA* is unable to sporulate. Top plates show the *S. venezuelae* wild-type (WT) growing on MYM agar for 1, 2, 3 and 4 days as indicated. Brown colouring indicates vegetative hyphal growth, white colouring indicates aerial hyphae formation and green colouring indicates sporulation because the mature spores are green. Bottom plates show cultures of the isogenic *ΔorrA* mutant grown for the same incubation times as indicated.

Vegetative colonies are stable, but emergent aerial hyphae yield genetically and phenotypically unstable mutants (Figure S1).

### Identification of OrrA target genes in *S. venezuelae*

To identify OrrA binding sites on the *S. venezuelae* genome the *ΔorrA* mutant was complemented by integrating a construct encoding OrrA with an N-terminal 3xFLAG tag *in trans* under the control of its native promoter. ChIP-seq analysis was performed in triplicate using this FLAG-tagged strain and wild-type *S. venezuelae* was used as a negative control. The sequencing data were analysed as described previously (23) and led to the identification of five promoters that were enriched in the FLAG-tagged strain relative to the control (Figure S2), including the *wblA* promoter (Figure 2) and the *vnz_04640* promoter (Figure S3).

**Figure 2.**
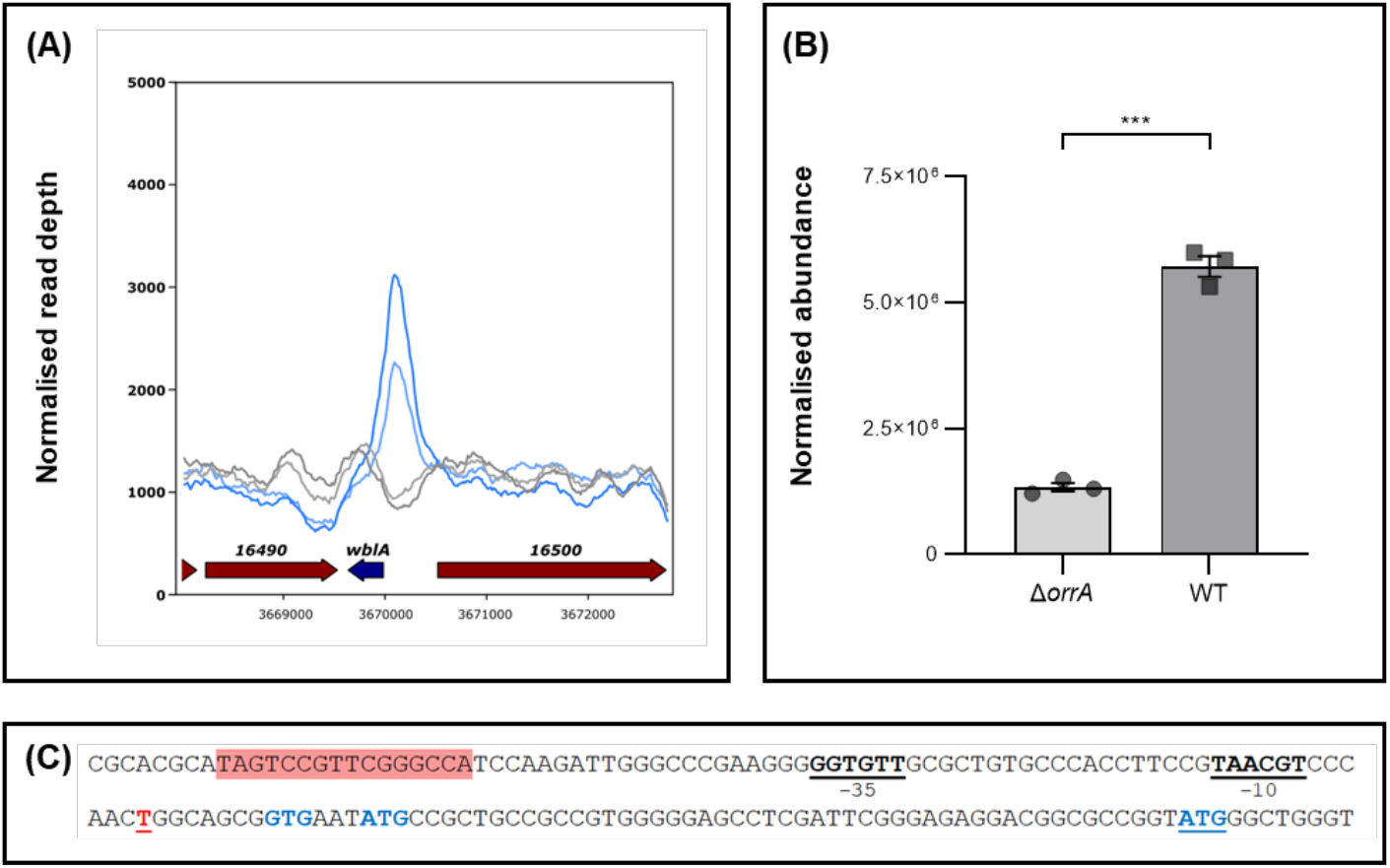
OrrA activates the production of WblA in *Streptomyces venezuelae*. (A). Normalised ChIP-seq read depth, blue traces show the OrrA-FLAG strain (n=2) and black traces show the wild-type control (n=2). (B). TMT proteomics data (n=3) showing the normalized peptide abundance of WblA in the *S. venezuelae* Δ*orrA* and wild-type strains at 18 hours growth on MYM medium. Individual replicate abundances are shown. *** = BH-adjusted *p*-value < 1 x 10^-7^ (background-based *t-*test). (C). The *wblA* promoter with OrrA consensus site highlighted in red, the RNA polymerase -35 and -10 sites underlined in black boldface, and the transcription start site (TSS) underlined in red boldface. There are three possible WblA translational start codons that are in frame, all marked in blue with the annotated start codon underlined. The same data are shown for OrrA target gene *vnz_04640* in Figure S3.

Four of these five promoters were used to measure OrrA binding *in vitro* using a DNA footprinting technique known as Reuseable DNA capture surface plasmon resonance, or ReDCaT SPR (Figures 3 and S4). ReDCaT SPR employs immobilised double-stranded oligonucleotide probes that span the promoter of interest. These probes are covalently attached to an SPR chip and subsequently challenged with the purified protein of interest (30). The full length OrrA protein could not be produced in soluble form in *E. coli*, so we used the purified DNA binding domain of OrrA for these experiments. This purified protein bound specifically to single probes at 4 of the 5 target promoters (Table 1). The *vnz_19230* promoter was not tested in these experiments because the ChIP-seq peak was overlooked in the initial analysis of the ChIP-seq data. It was identified as an OrrA target after scanning the *S. venezuelae* genome with the experimentally verified OrrA consensus binding sequence.

**Table 1.**
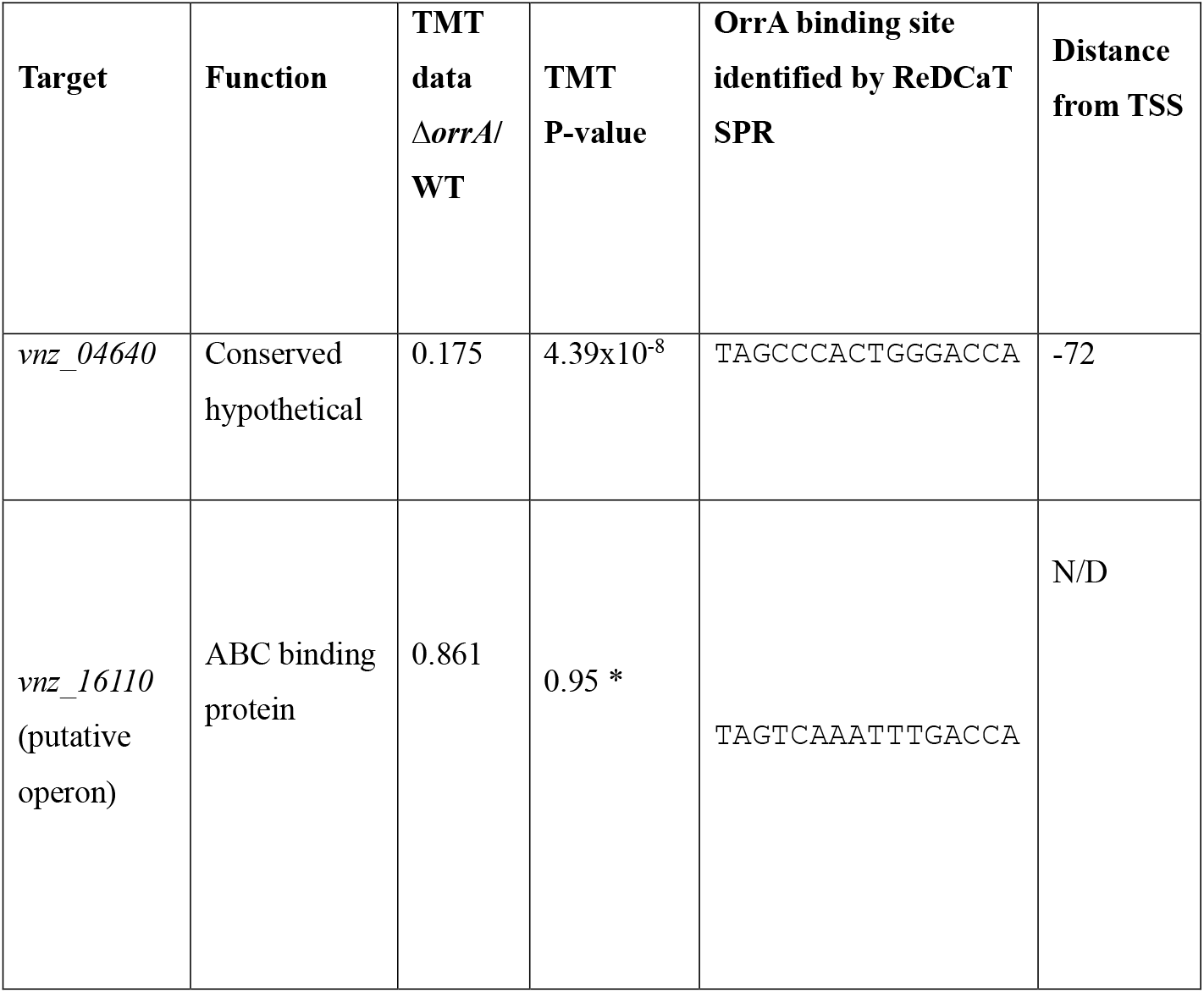
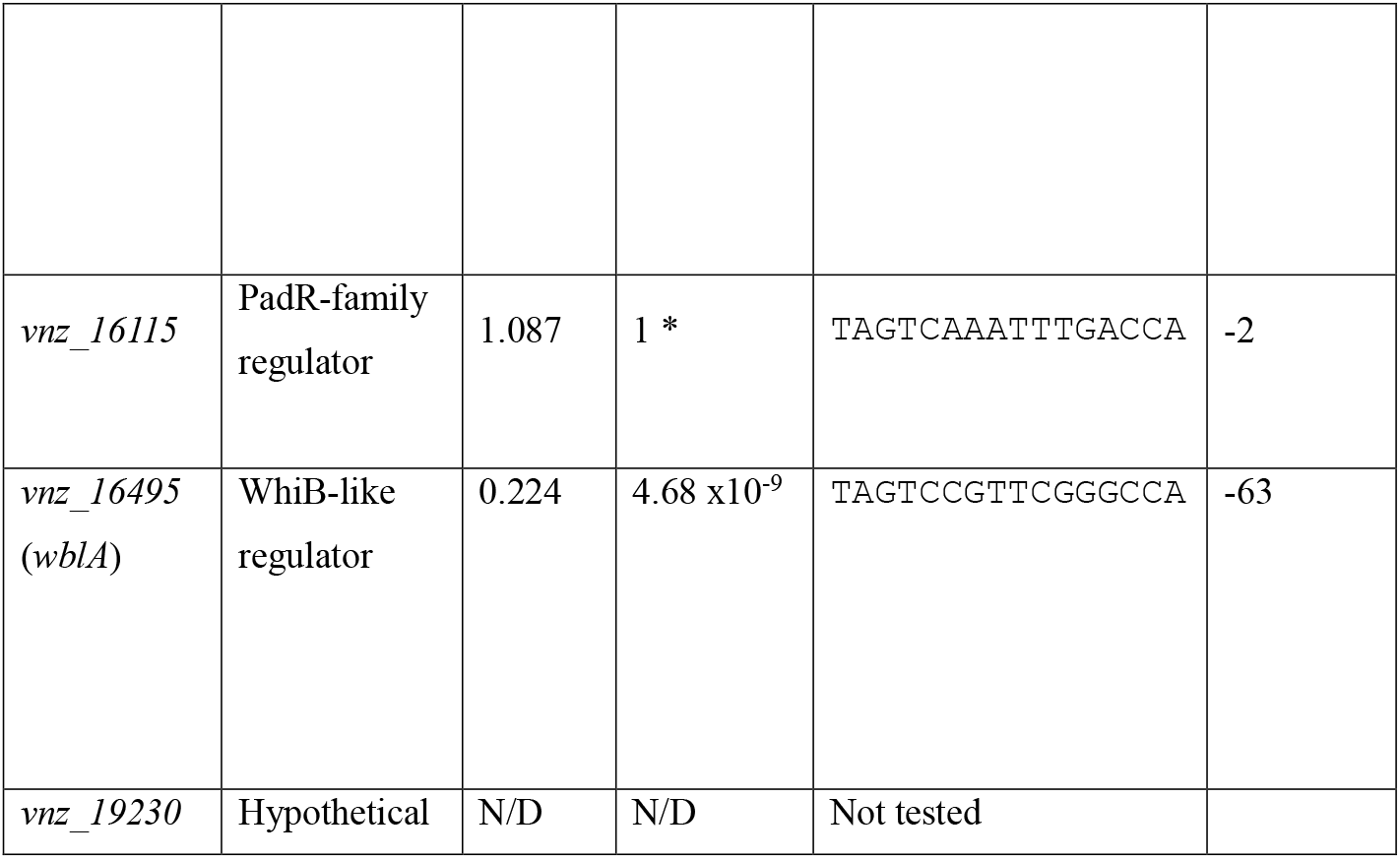
OrrA binding sites in *S. venezuelae*. ChIP-seq against FLAG-tagged OrrA enriched these gene promoters (Figure S2) and the OrrA binding sites were identified for four of them *in vitro* using ReDCaT SPR. TMT proteomics data is shown for 18-hour cultures and indicate that OrrA activates expression of *wblA* and *vnz_04640* which encodes a conserved small hypothetical protein. N/D = not detected. *P-value not significant (>0.05).

**Figure 3.**
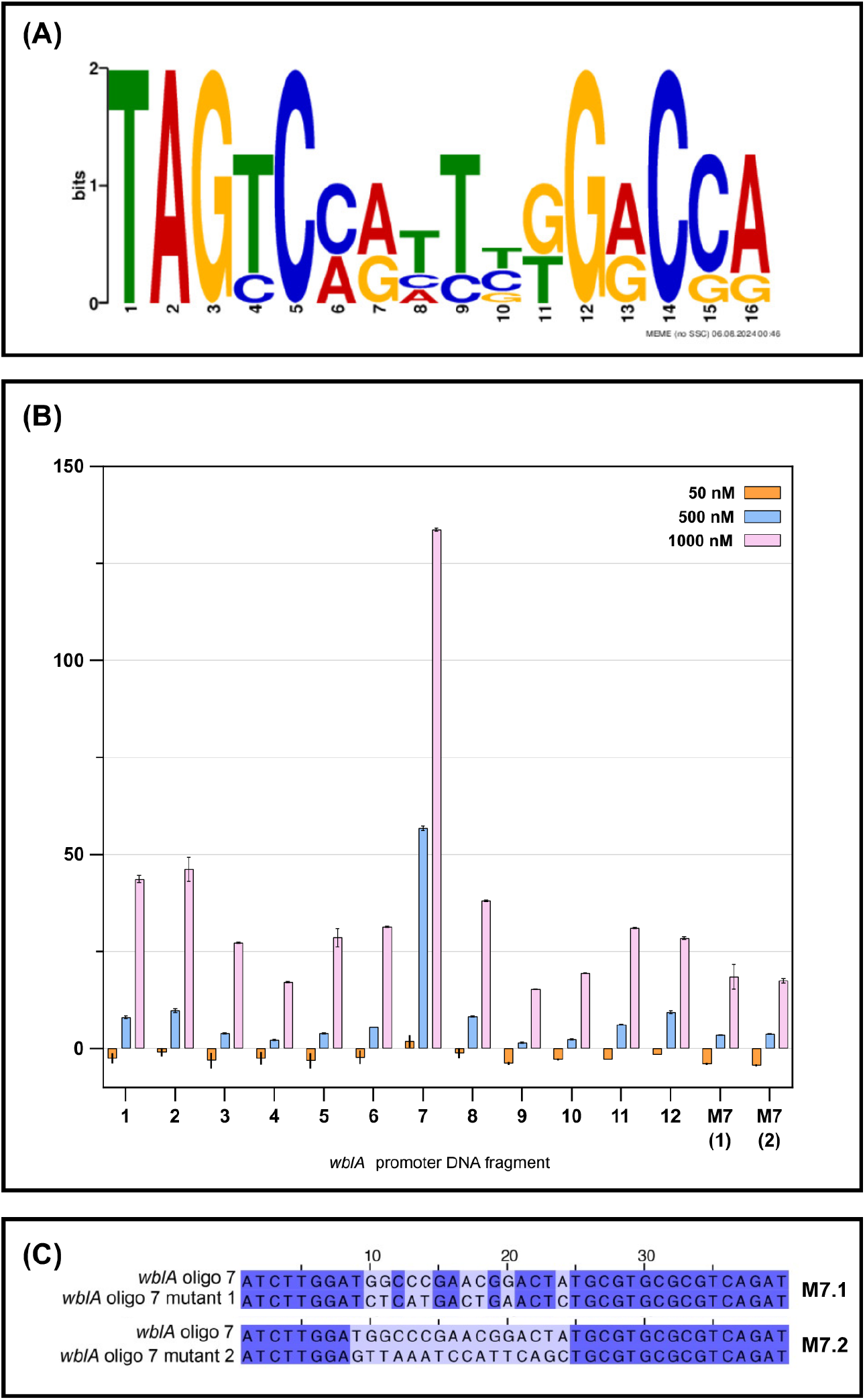
Identification of the OrrA binding site. (A). MEME generated consensus binding site for OrrA generated by aligning the four OrrA binding sites that were experimentally verified using ReDCaT SPR (Figure S4). (B). Binding of OrrA-DBD to the double stranded fragments of *wblA* promoter DNA was determined via ReDCaT SPR. Binding strength is represented as %R_Max_ values, calculated assuming two OrrA-DBDs binding to the imperfect palindromic MEME consensus; error bars shown (± SEM, n = 2). The key shows the concentrations of oligonucleotides used in SPR experiments. OrrA binds specifically to oligo probe 7 and mutation of the OrrA consensus site on probe 7 (M7.1 and M7.2) reduced OrrA binding to background levels. (C). Alignment of wild-type probe 7 with mutated probes M7.1 (partially mutated) and M7.2 (fully mutated). Note that the consensus is the reverse complement of the MEME consensus in A.

Alignment of the OrrA binding sites identified using ReDCaT SPR enabled an OrrA consensus DNA recognition sequence to be generated using MEME (31) and this was validated by mutating conserved nucleotides in the ReDCaT SPR probes (Figures 3 and S4). The binding site appears to be palindromic which is consistent with dimeric OrrA binding to its target promoters.

Tandem-mass-tag (TMT) proteomics analysis of the wild-type and *ΔorrA* strains grown for 18 hours on MYM agar show that the levels of WblA and Vnz_04640 are reduced in the *ΔorrA* mutant relative to wild-type (Figures 2 and S3 and Table 1; see Table S5 for the complete proteomics dataset). We conclude that OrrA directly activates the production of both proteins, which agrees with an earlier study that identified these genes as targets for *S. coelicolor* OrrA (13). Vnz_04640 is a small hypothetical 67 amino acid protein that is highly conserved in *Streptomyces* species and restricted to the phylum Actinomycetota (Figure S5). A previous study reported that over-expression of either *wblA* or the *vnz_04640* homologue *sco1375* in *S. coelicolor* only partially restores development in an *S. coelicolor ΔorrA* mutant (13). To test this in *S. venezuelae*, we over-expressed *vnz_04640* and *wblA* individually and together in the *ΔorrA* mutant. Over-expression of *vnz_04640* had no visible effect on the growth and development of either the wild-type or *ΔorrA* strains (Figure 4A). However, over-expression of either *wblA* or an artificial operon of *vnz_04640-wblA* fully restored sporulation to the *ΔorrA* mutant suggesting the *ΔorrA* developmental phenotype is due to the reduction in the levels of WblA (Figures 4B and C).

**Figure 4.**
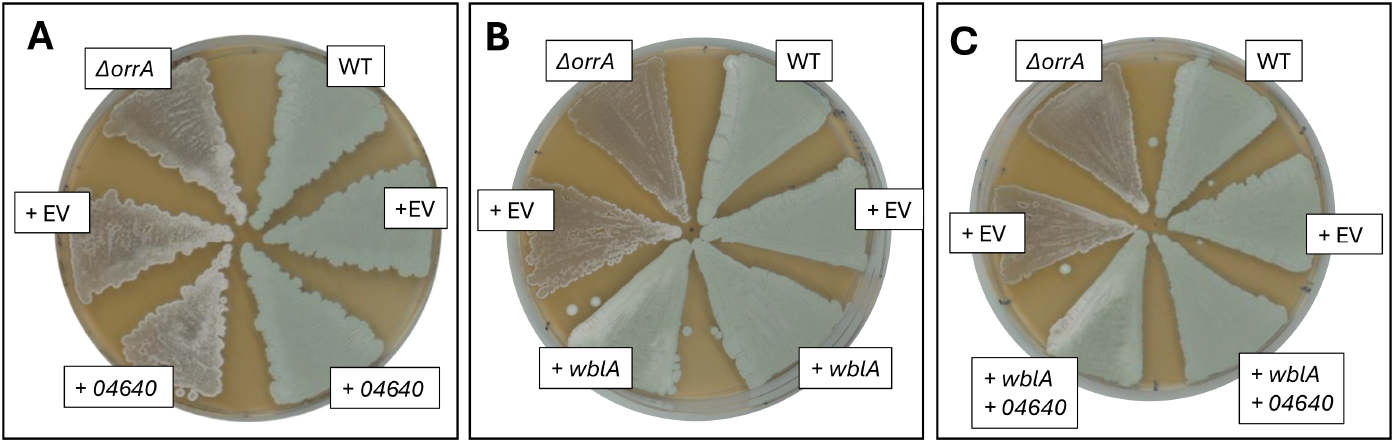
Over-expression of *wblA* restores normal sporulation to the *ΔorrA* mutant. (A). Wild-type and *ΔorrA* strains complemented in trans with either empty pIJ10257 (+EV) or pIJ10257 containing *vnz_04640* downstream of the *ermE\** promoter. Cultures were grown for 3 days at 30 °C on MYM agar.

ChIP-seq and ReDCaT SPR data indicate that OrrA also binds upstream of the *vnz_16105-10* operon encoding putative drug resistance transporters and the *vnz_16115* gene which encodes a putative transcription factor that is a member of the σ^E^ regulon in *S. coelicolor* (Table 1) (32). Of these three genes, products were only detected in the TMT proteomics experiment for *vnz_16110* and *vnz_16115* and neither were significantly affected by the loss of OrrA (Table 1).

### *S. venezuelae ΔorrA* and *ΔwblA* are defective in sporulation and over produce chloramphenicol

The data in Figure 4 show that over-expression of *wblA* rescues the *ΔorrA* mutant and restores normal growth and sporulation suggesting a reduction in the levels of WblA results in the *ΔorrA* phenotype. To test this further in *S. venezuelae*, we deleted the *wblA* gene and observed that the mutant phenotype exhibits similar growth and development to the *ΔorrA* mutant in that it does not sporulate (Figure 5).

**Figure 5.**
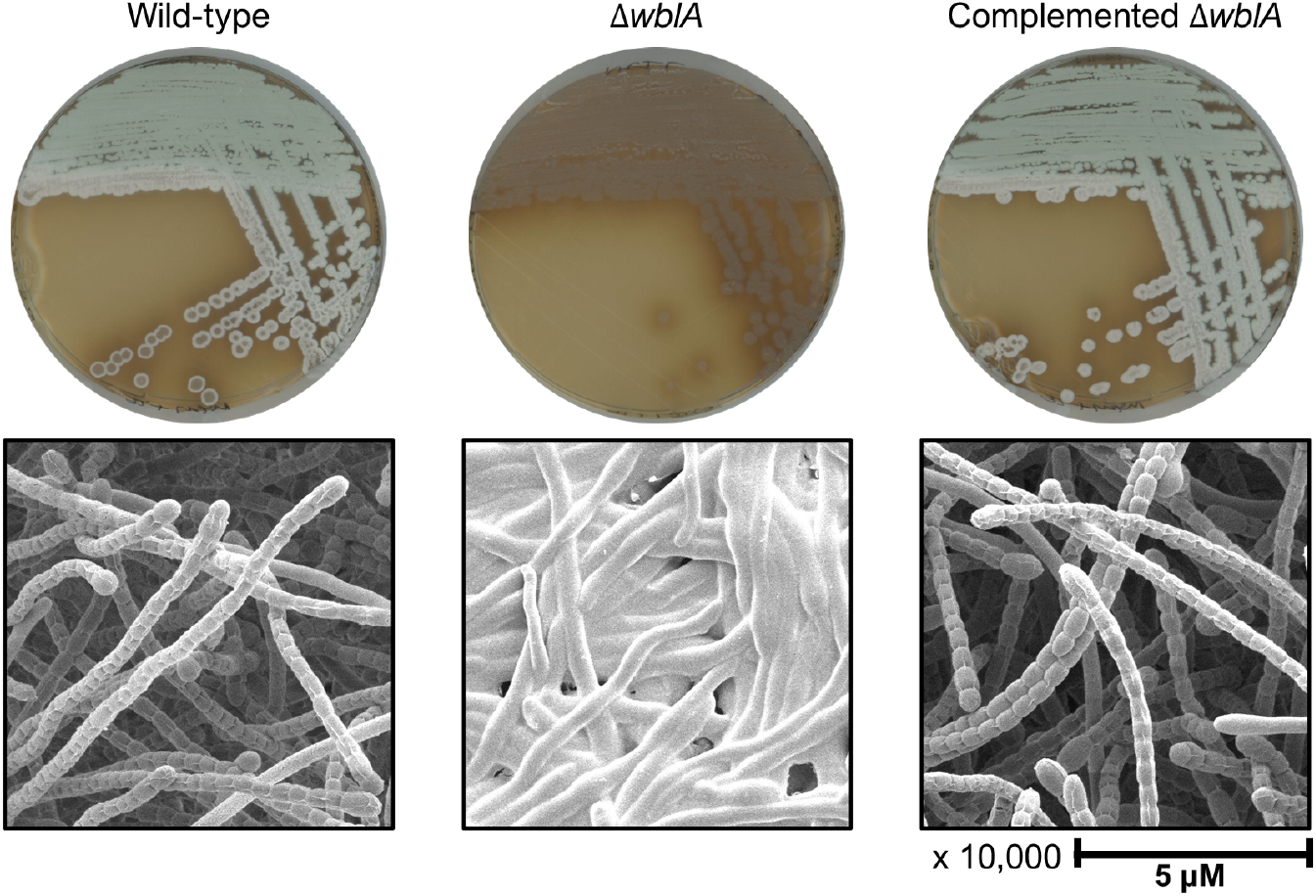
*S. venezuelae ΔwblA* does not sporulate. Top panels show MYM agar plate grown cultures of the wild-type (WT) and isogenic *ΔwblA* and *ΔwblA* complemented strains grown for three days. *S. venezuelae* spores are green, the WT and *ΔwblA* complemented strain are sporulating, whereas the *ΔwblA* mutant is not forming spores. Bottom panels show scanning electron micrographs of the same strains and confirm that the WT and *ΔwblA* complemented strain form aerial hyphae and spores, while the *ΔwblA* strain is only forming vegetative substrate mycelium.

Given that *S. coelicolor ΔorrA* and *ΔwblA* mutants both overproduce antibiotics (13), it was hypothesised that *S. venezuelae ΔorrA* would do the same. *S. venezuelae* makes the antibiotic chloramphenicol (33) and TMT proteomics data comparing the *S. venezuelae* wild-type and *ΔorrA* strains demonstrated that the entire chloramphenicol biosynthetic pathway is significantly upregulated in the *orrA* mutant (Figure 6A). We tested the antibacterial activities of *S. venezuelae* wild-type, *ΔorrA* and *ΔwblA* strains using agar plate bioassays. The results show that the wild-type strain does not inhibit *B. subtilis* while both mutants have clear zones of inhibition in overlay bioassays (Figure 6B). Next, we extracted cultures of the wild-type, *ΔorrA* and *ΔwblA* strains and showed by LCMS analysis that chloramphenicol is produced by both mutant strains but not by wild-type *S. venezuelae* (Figure 6C). These data indicate that the reduction in WblA levels in the *ΔorrA* strain is responsible for activating chloramphenicol biosynthesis.

**Figure 6.**
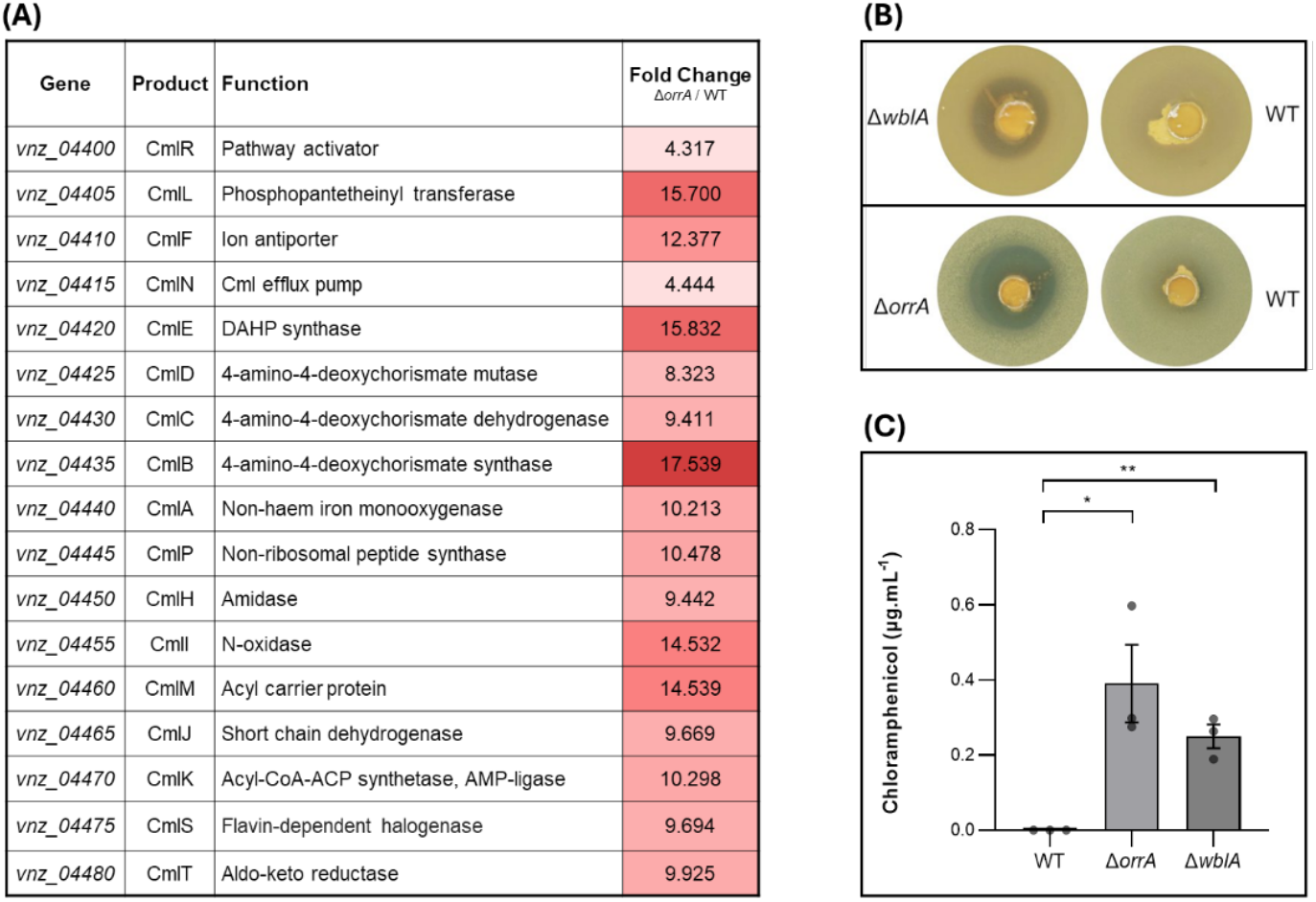
Chloramphenicol biosynthesis is upregulated in the *ΔorrA* mutant. (A). Significant fold changes in peptide abundance observed for the proteins encoded by the chloramphenicol BGC in the *ΔorrA* mutant relative to wild-type (WT) strains, quantified by TMT proteomics. (B). Agar plate bioassays showing colonies of wild-type S. venezuelae (WT) and the Δ*wblA* and Δ*orrA* mutants overlaid with soft agar containing the indicator strain *Bacillus subtilis*. The mutants inhibit *B. subtilis* but WT does not. (C). LCMS measurements of extracted cultures show that the Δ*orrA* and Δ*wblA* mutants are both producing the antibiotic chloramphenicol (n=3). *p*-values, * = 0.0198, ** *p =* 0.0140 (Unpaired *t*-test).

## 7. Concluding Remarks

In this work we identified two genes under the direct control of the orphan response regulator OrrA in *S. venezuelae* NRRL B-65442, namely *wblA* and *vnz_04640*. We also showed that the *ΔorrA* and *ΔwblA* strains are phenotypically very similar, with an inability to sporulate and overproduction of the antibiotic chloramphenicol. Over-expression of *wblA* in the *ΔorrA* strain restored normal sporulation suggesting that the *ΔorrA* developmental phenotype is primarily due to a reduction in the cellular levels of WblA. We found that the *ΔwblA* phenotype is stable whereas the *ΔorrA* phenotype can revert to a strain that forms aerial hyphae (Figure S1).

A study of *S. coelicolor* also identified *wblA* and *sco1375* (the *vnz_04640* homologue) as OrrA target genes and we propose that OrrA function is highly conserved in the genus *Streptomyces*. Overexpression of *sco1375* or *wblA* partially restored normal development to *S. coelicolor ΔorrA* whereas overexpression of *vnz_04640* has no effect on the *ΔorrA* mutant and overexpression of *wblA* restored sporulation (Figure 4). WblA is required for normal aerial hyphae formation and sporulation (13,14) and it is a global regulator of antibiotic biosynthesis in *Streptomyces* species (19). Consistent with this, the deletion of *S. coelicolor orrA* leads to a reduction in WblA and an increase in actinorhodin and undecylprodigiosin biosynthesis in *S. coelicolor* while deletion of *S. venezuelae ΔorrA* reduces WblA levels and increases chloramphenicol biosynthesis (Figures 2 and 6).

The *sco1375* and *vnz_04640* genes encode small hypothetical proteins that contain 67 and 68 amino acids, respectively, and both contain DUF5999, a domain of unknown function which is predominantly found in the phylum Actinomycetota. The function of this protein is not known but it is highly conserved in *Streptomyces* species, as are OrrA and WblA, suggesting they have conserved functions (Figure S5), with OrrA solely dedicated to activating the production of Vnz_04640 and WblA (Figures 2 and S3). Future work will be aimed at investigating the functions of Vnz_04640 and WblA in *S. venezuelae*.

## Supporting information

TMT proteomics data

## 8. Author statements

### 8.1 Author contributions

Conceptualisation: KRT, LB, NAH, BW and MIH

Data curation: KRT, LB, NAH, ADMB, CA, GC, GS, RS, BW and MIH

Formal analysis: KRT, LB, NAH, ADMB, GC, GS

Funding acquisition: BW and MIH

Investigation: KRT, NAH, ADMB, CA

Methodology: KRT, NAH, LB, CA RS, GC, GS

Project administration: BW, MIH

Resources: RS, GS, BW, MIH

Software: GC, GS

Supervision: LB, BW and MIH

Visualisation: KRT

Writing – original draft: KRT and MIH

Writing – review and editing: all authors.

### 8.2 Conflicts of interest

The authors declare that there are no conflicts of interest.

### 8.3 Funding information

Kelly-Rose Tulley, Lucas Balis and Courtney Andrews: BBSRC doctoral training programme grant BB/M011216/1. Ainsley D. M. Beaton: BBSRC responsive mode grant BB/W000628/1. Neil Holmes, Govind Chandra, Gerhard Saalbach and Rhea Stringer: Institute Strategic Programme Project BBS/E/J/000PR9790. The John Innes Centre Bioimaging Platform is supported by the UKRI Biotechnology and Biological Sciences Research Council (grant BB/CCG2240/1).

### 8.4 Ethical approval

N/A

### 8.5 Consent for publication

N/A

## 8.6 Acknowledgements

We thank present and past members of the Hutchings and Wilkinson research groups and our colleagues in the department for Molecular Microbiology for helpful discussions.

## Notes

### Competing Interest Statement

The authors have declared no competing interest.

## References

1. Schlimpert S, Elliot MA. The Best of Both Worlds—Streptomyces coelicolor and Streptomyces venezuelae as Model Species for Studying Antibiotic Production and Bacterial Multicellular Development. J Bacteriol. 2023 Jun 22;e00153–23.

2. Hutchings MI, Truman AW, Wilkinson B. Antibiotics: past, present and future. Curr Opin Microbiol. 2019 Oct;51:72–80.

3. Bush MJ, Tschowri N, Schlimpert S, Flärdh K, Buttner MJ. c-di-GMP signalling and the regulation of developmental transitions in streptomycetes. Nat Rev Microbiol. 2015 Dec;13(12):749–60.

4. Cruz-Bautista R, Ruíz-Villafán B, Romero-Rodríguez A, Rodríguez-Sanoja R, Sánchez S. Trends in the two-component system’s role in the synthesis of antibiotics by Streptomyces. Appl Microbiol Biotechnol. 2023 Aug;107(15):4727–43.

5. McLean TC, Lo R, Tschowri N, Hoskisson PA, Al Bassam MM, Hutchings MI, et al. Sensing and responding to diverse extracellular signals: an updated analysis of the sensor kinases and response regulators of Streptomyces species. Microbiology. 2019 Sep 1;165(9):929–52.

6. Som NF, Heine D, Holmes NA, Munnoch JT, Chandra G, Seipke RF, et al. The Conserved Actinobacterial Two-Component System MtrAB Coordinates Chloramphenicol Production with Sporulation in Streptomyces venezuelae NRRL B-65442. Front Microbiol. 2017 Jun 28;8:1145.

7. Som NF, Heine D, Holmes N, Knowles F, Chandra G, Seipke RF, et al. The MtrAB two-component system controls antibiotic production in Streptomyces coelicolor A3(2). Microbiology. 2017 Oct 1;163(10):1415–9.

8. Tian J, Li Y, Zhang C, Su J, Lu W. Characterization of a pleiotropic regulator MtrA in Streptomyces avermitilis controlling avermectin production and morphological differentiation. Microb Cell Factories. 2024 Apr 8;23(1):103.

9. Zhu Y, Zhang P, Zhang J, Wang J, Lu Y, Pang X. Impact on Multiple Antibiotic Pathways Reveals MtrA as a Master Regulator of Antibiotic Production in Streptomyces spp. and Potentially in Other Actinobacteria. Appl Environ Microbiol. 2020 Oct;86(20):e01201–20.

10. Hoskisson PA, Hutchings MI. MtrAB–LpqB: a conserved three-component system in actinobacteria? Trends Microbiol. 2006 Oct;14(10):444–9.

11. Beaton ADM, Devine R, Holmes NA, Som NF, Balis L, Noble K, et al. MtrAB activates ectoine production and triggers sporulation in response to osmotic stress in Streptomyces venezuelae. bioRxiv 10.64898/2025.12.19.695424.

12. Zheng G, Liu P, He W, Tao H, Yang Z, Sun C, et al. Identification of the cognate response regulator of the orphan histidine kinase OhkA involved in both secondary metabolism and morphological differentiation in Streptomyces coelicolor. Appl Microbiol Biotechnol. 2021 Aug;105(14–15):5905–14.

13. Zhu Y, Wang X, Zhang J, Ni X, Zhang X, Tao M, et al. The regulatory gene wblA is a target of the orphan response regulator OrrA in Streptomyces coelicolor. Environ Microbiol. 2022 Apr 10;1462-2920.15992.

14. Fowler-Goldsworthy K, Gust B, Mouz S, Chandra G, Findlay KC, Chater KF. The actinobacteria-specific gene wblA controls major developmental transitions in Streptomyces coelicolor A3(2). Microbiology. 2011 May 1;157(5):1312–28.

15. Bush MJ. The actinobacterial WhiB-like (Wbl) family of transcription factors: The Actinobacterial WhiB-like (Wbl) Family of Transcription Factors. Mol Microbiol. 2018 Dec;110(5):663–76.

16. Guiza Beltran D, Wan T, Zhang L. WhiB-like proteins: Diversity of structure, function and mechanism. Biochim Biophys Acta BBA - Mol Cell Res. 2024 Oct;1871(7):119787.

17. Jordan ML, Schlimpert S. Microbe Profile: Streptomyces venezuelae – a model species to study morphology and differentiation in filamentous bacteria: Microbiology 2025 Mar 13;171(3).

18. Gomez-Escribano JP, Holmes NA, Schlimpert S, Bibb MJ, Chandra G, Wilkinson B, et al. Streptomyces venezuelae NRRL B-65442: genome sequence of a model strain used to study morphological differentiation in filamentous actinobacteria. J Ind Microbiol Biotechnol. 2021 Dec 23;48(9–10):kuab035.

19. Nah HJ, Park J, Choi S, Kim ES. WblA, a global regulator of antibiotic biosynthesis in Streptomyces. J Ind Microbiol Biotechnol. 2021 Jun 4;48(3–4):kuab007.

20. Kiester T, Bibb MJ, Buttner MJ, Chater KF and Hopwood DA. Practical Streptomyces Genetics [Internet]. Norwich: The John Innes Foundation; 2000. Available from: https://streptomyces.org.uk/PracticalStreptomycesGenetics.pdf

21. Cobb RE, Wang Y, Zhao H. High-Efficiency Multiplex Genome Editing of Streptomyces Species Using an Engineered CRISPR/Cas System. ACS Synth Biol. 2015 Jun 19;4(6):723– 8.

22. Feeney MA, Newitt JT, Addington E, Algora-Gallardo L, Allan C, Balis L, et al. ActinoBase: tools and protocols for researchers working on Streptomyces and other filamentous actinobacteria. Microb Genomics. 2022 Jan 1;8(7):mgen000824.

23. Devine R, McDonald HP, Qin Z, Arnold CJ, Noble K, Chandra G, et al. Re-wiring the regulation of the formicamycin biosynthetic gene cluster to enable the development of promising antibacterial compounds. Cell Chem Biol. 2021 Apr;28(4):515-523.e5.

24. Devine R, Noble K, Stevenson C, De Oliveira Martins C, Saalbach G, McDonald HP, et al. Redox control of antibiotic biosynthesis. mBio. 2025. 16:e01369–25.

25. Wessel D, Flügge UI. A method for the quantitative recovery of protein in dilute solution in the presence of detergents and lipids. Anal Biochem. 1984 Apr 1;138(1):141–3.

26. McLean TC, Beaton ADM, Martins C, Saalbach G, Chandra G, Wilkinson B, et al. Evidence of a role for CutRS and actinorhodin in the secretion stress response in Streptomyces coelicolor M145. Microbiology [Internet]. 2023 Jul 7;169(7).

27. Katoh K, Standley DM. MAFFT Multiple Sequence Alignment Software Version 7: Improvements in Performance and Usability. Mol Biol Evol. 2013 Apr 1;30(4):772–80.

28. Afgan E, Baker D, Batut B, van den Beek M, Bouvier D, Čech M, et al. The Galaxy platform for accessible, reproducible and collaborative biomedical analyses: 2018 update. Nucleic Acids Res. 2018 Jul 2;46(W1):W537–44.

29. Letunic I, Bork P. Interactive Tree Of Life (iTOL) v5: an online tool for phylogenetic tree display and annotation. Nucleic Acids Res. 2021 Jul 2;49(W1):W293–6.

30. Stevenson CEM, Lawson DM. Analysis of Protein–DNA Interactions Using Surface Plasmon Resonance and a ReDCaT Chip. In: Daviter T, Johnson CM, McLaughlin SH, Williams MA, editors. Protein-Ligand Interactions: Methods and Applications. New York, NY: Springer US; 2021. p. 369–79.

31. Bailey TL, Johnson J, Grant CE, Noble WS. The MEME Suite. Nucleic Acids Res. 2015 Jul 1;43(W1):W39–49.

32. Tran NT, Huang X, Hong H, Bush MJ, Chandra G, Pinto D, et al. Defining the regulon of genes controlled by σ E, a key regulator of the cell envelope stress response in Streptomyces coelicolor. Mol Microbiol. 2019 Aug;112(2):461–81.

33. Fernández-Martínez LT, Borsetto C, Gomez-Escribano JP, Bibb MJ, Al-Bassam MM, Chandra G, et al. New Insights into Chloramphenicol Biosynthesis in Streptomyces venezuelae ATCC 10712. Antimicrob Agents Chemother. 2014 Dec;58(12):7441–50.

