## Supplementary Tables, Figures and Methods for "Defining the OrrA regulon and its role in development and antibiotic production in *Streptomyces venezuelae* NRRL B-65442"

**SUPPLEMENTARY DATA**

**Table S1. Strains used in this study**

| ***Escherichia coli* strains** | | | |
|---|---|---|---|
| **Strain name** | **Description** | **Source** | **Reference** |
| BL21 (DE3) | fhuA2 [lon] ompT gal (λ DE3) [dcm] ΔhsdSλ DE3 = λsBamHIo ΔEcoRI-B | New England Biolabs | |
| DH5α | fhuA2 D (argF-lacZ)U169 IphoA glnV44 Φ80lacZ DM15 gyrA96 recA1 relA1 endA1 thi-1 | New England Biolabs | |
| ET12567 + pUZ8002 | Methylation deficient dcm-dam- strain + driver plasmid used for conjugation into *Streptomyces* species | John Innes Centre http://streptomyces.org.uk | |
| ***Streptomyces venezuelae* NRRL B-65442 strains** | | | |
| **Strain name** | **Description** | **Source** | **Reference** |
| *S. venezuelae* NRRL B-65442 | Model wild-type strain, forms green spores, sporulates in solid and liquid culture | John Innes Centre http://streptomyces.org.uk | Gomez-Escribano *et al.*, 2021; Schlimpert and Elliot, 2023; Jordan and Schlimpert, 2025) |

| | | | |
|---|---|---|---|
| Δ*orrA* | In-frame deletion of the *orrA* gene | This work | This work |
| Δ*orrA* + *orrA* Flag | Δ*orrA* complemented *in trans* with a construct encoding 3xFlag-OrrA under its native promoter. Used for ChIP-seq | This work | This work |
| Δ*orrA* + *ermEp**-*vnz_04640* | Δ*orrA* complemented *in trans* with a pIJ10257 construct expressing *vnz_04640* from the *ermE** promoter | This work | This work |
| Δ*orrA* + *ermEp**-*wblA* | Δ*orrA* complemented *in trans* with a pIJ10257 construct expressing *wblA* from the *ermE** promoter | This work | This work |
| Δ*orrA* + *ermEp**-*vnz_04640-wblA* | Δ*orrA* complemented *in trans* with a pIJ10257 construct expressing an artificial operon of *vnz_04640-wblA* from the *ermE** promoter | This work | This work |
| Δ*wblA* | In-frame deletion of the *wblA* gene | This work | This work |
| Δ*wblA* + *wblA* | Δ*wblA* complemented *in trans* with *wblA* under its native promoter | This work | This work |

**Table S2. Plasmids used in this study.**

| Plasmid | Description | Origin | Resistance marker |
|---|---|---|---|
| pIJ10257 | Integrative vector based on pMS81 for the over-expression of genes using the high-level constitutive *ermEp** promoter. oriT, ΦBT1 attB-int, HygR, *ermEp**, pMS81 backbone | John Innes Centre, http://streptomyces.org.uk (Hong *et al.*, 2005) | Hyg |
| pCRISPomyces-2 | AprR, oriT, reppSG5(ts) , oriColE1, sSpcas9, synthetic guide RNA cassette | (Cobb, Wang and Zhao, 2015) | Apr |
| pCRISPomyces-2 *orrA* | For in-frame deletion of *S. venezuelae orrA* | This work | Apr |
| pCRISPomyces-2 *wblA* | For in-frame deletion of *S. venezuelae wblA* | This work | Apr |
| pIJ10257-*orrA* | *orrA* over-expression vector | This work | Hyg |
| pIJ10257-*vnz_04640* | *vnz_04640* over-expression vector | This work | Hyg |
| pIJ10257-*wblA* | *wblA* over-expression vector | This work | Hyg |
| pIJ10257-*vnz_04640-wblA* | *vnz_04640-wblA* over-expression vector | This work | Hyg |
| pET29a | pBR322 origin and fI origin, KanR, expression vector | Invitrogen | Kan |
| pET29a-*orrA* DBD | For *E. coli* over-production and purification of the OrrA DNA binding domain | This work | Kan |

| | | | |
|---|---|---|---|
| | (DBD) using nickel affinity chromatography | | |
| pIJ10770 (aka pSS170) | Phage-based ΦBT1 integrative vector derived from pMS82. oriT, ΦBT1 attB-int, HygR, derived from pMS82 | John Innes Centre<br><br>(Schlimpert *et al.*, 2017)<br><br>30/01/2026 13:44:00 | Hyg |
| pIJ10770 + *wblA* | For *in trans* complementation of *S. venezuelae* Δ*wblA* | This work | Hyg |
| pSS170 + *orrA* N-FLAG | *In trans* production of N-terminally Flag-tagged OrrA for ChIP-seq. | This work | Hyg |
| pSS170 *wblA* long | For *in trans* complementation of the *S. venezuelae* Δ*wblA* mutant | This work | Hyg |
| pSS170 *orrA* | For *in trans* complementation of the *S. venezuelae* Δ*orrA* mutant | This work | Hyg |

**Table S3. Primers used in this study.**

| Primer Description | Sequence |
|---|---|
| pCRISP *orrA* homologous region 1 - Forward | GCTCGGTTGCCGCCGGGCGTTTTTATCTAGAGTTGCGCCTACAGGACAGGA |
| pCRISP *orrA* homologous region | GCTGCTGCGACCAGGCGAGCTCGCGAAGCTGTCCGCCATCGTTC |

| | |
|---|---|
| 1 - Reverse | |
| pCRISP *orrA* homologous region 2 - Forward | GCGAGCTCGCCTGGTCGCAGCAGCCTGGAGATCCGCTAGCGGCC |
| pCRISP *orrA* homologous region 2 - Reverse | GCAACGCGGCCTTTTTACGGTTCCTGGCCTCTAGACCAGGTCCAGCTGGACACG |
| *orrA* deletion test - Forward 1 | GCGGCTGTGATCGCATCCTCTCCC |
| *orrA* deletion test - Forward 2 | CCCTGAGTAGCGGTTGGAGGACCG |
| *orrA* deletion test - Reverse 1 | CCGGAGCTGACGAAGCCCCTGG |
| *orrA* deletion test - Reverse 2 | CCGAACTGTCCGCCGACGAAGCC |
| *orrA* gRNA Forward | ACGCCTGGATCATCGACTTGAACT |

| | |
|---|---|
| *orrA* gRNA Reverse | AAACAGTTCAAGTCGATGATCCAG |
| pET29a *orrA* DBD C-terminal 6xHis - Forward | CTTTAAGAAGGAGATATACATATGGTGCCCGCGCCCCGGCTG |
| pET29a *orrA* DBD C-terminal 6xHis - Reverse | GTGCTCGAGTGCGGCCGCAAGCTTGCGGATCTCCAGGATCTTCTCCCG |
| pET29a *orrA* DBD N-terminal 6xHis - Forward | CTGGTGCCGCGCGGCAGCCATATGGTGCCCGCGCCCCGGCTG |
| pET29a *orrA* DBD N-terminal 6xHis - Reverse | TCGAGTGCGGCCGCAAGCTTCTAGCGGATCTCCAGGATCTTCTCCCG |
| pCRISP *wblA* homologous region 1 - Forward | GCTCGGTTGCCGCCGGGCGTTTTTATCTAGATGGCCTGCGCGACCTTCTCG |
| pCRISP *wblA* homologous region 1 - Reverse | GCTGCTGCGACCAGGCGAGCTCGCACCGGCGCCGTCCTCTCC |

| | |
|---|---|
| pCRISP *wblA* homologous region 2 - Forward | GCGAGCTCGCCTGGTCGCAGCAGCCACCCGTCCGGCGGCTGA |
| pCRISP *wblA* homologous region 2 - Reverse | GCAACGCGGCCTTTTTACGGTTCCTGGCCTCTAGAGGCGATGGAGAAGCTGGGCCA |
| *wblA* gRNA Forward | ACGCCCGGATGAACTTTTCGTACA |
| *wblA* gRNA Reverse | AAACTGTACGAAAAGTTCATCCGG |
| pSS170 *wblA* - Forward | CAGCAAAAGGGGATGATAAGTTTATCAAGCTTCTCATTCGCCGGACACACGTATATG |
| pSS170 *wblA* - Reverse | CTTAATTAACTCGAGAGATGTACACCTAGGCTAGCCGACCGCGGCGTA |
| pIJ10257 *wblA* - Forward | GTCTAGAACAGGAGGCCCCATATGATG CCG CTG CCG CCG T |
| pIJ10257 *wblA* - Reverse | CATGAGAACCTAGGATCCAAGCTTCTAGCCGACCGCGGCGTA |
| *wblA* test (within HRT) - Forward | CCTAAGCTGCTCCCAACTGTCAC |

| | |
|---|---|
| *wblA* test (within HRT) - Reverse | GGAAGACTGACCGAAGTGGCT |
| *wblA* test (outside HRT) - Forward | GCGCTTGGTCGCTTCCTGGAT |
| *wblA* test (outside HRT) - Reverse | CGAACTGCACGCGATGATGCT |
| pIJ10257 *vnz04640* - Forward | TCGTCTAGAACAGGAGGCCCCATATGTGCCAGCACCAGCCTGCCTG |
| pIJ10257 *vnz04640* - Reverse | CTCATGAGAACCTAGGATCCAAGCTTTCAGGCGGCGGTCGTCACCTG |
| pIJ10257 *vnz04640* *wblA* – Forward 1 | TCGTCTAGAACAGGAGGCCCCATATGCCGCTGCCGCCGTGGGGGAG |
| pIJ10257 *vnz04640* *wblA* – Forward 2 | CTCAACATCGGAGGTAAGCCATGTGCCAGCACCAGCCTGC |
| pIJ10257 *vnz04640* *wblA* – Reverse 1 | GGCTTACCTCCGATGTTGAGCTAGCCGACCGCGGCGTACG |
| pIJ10257 *vnz04640* *wblA* – Reverse 2 | ATGAGAACCCTAGGGGATCCAAGCTTTCAGGCGGCGGTCGTCACCTG |

**Table S1. ReDCaT SPR oligonucleotides used to make double stranded probes. Synthesised by Integrated DNA Technologies (IDT).**

| Label | Sequence | ds MW (da) |
|---|---|---|
| KR_SPR_wblA_1_F | GCCGCCTGCGCACTCCAGTCAGCTACCCAGCCCATACCGG | 30744.68 |
| KR_SPR_wblA_1_R | CCGGTATGGGCTGGGTAGCTGACTGGAGTGCGCAGGCGGCcctaccctacgtcctcctgc | |
| KR_SPR_wblA_2_F | CCCAGCCCATACCGGCGCCGTCCTCTCCCGAATCGAGGCT | 30744.68 |
| KR_SPR_wblA_2_R | AGCCTCGATTCGGGAGAGGACGGCGCCGGTATGGGCTGGGcctaccctacgtcctcctgc | |
| KR_SPR_wblA_3_F | TCCCGAATCGAGGCTCCCCCACGGCGGCAGCGGCATATTC | 30743.7 |
| KR_SPR_wblA_3_R | GAATATGCCGCTGCCGCCGTGGGGGAGCCTCGATTCGGGAcctaccctacgtcctcctgc | |
| KR_SPR_wblA_4_F | GGCAGCGGCATATTCACCGCTGCCAGTTGGGACGTTACGG | 30741.74 |
| KR_SPR_wblA_4_R | CCGTAACGTCCCAACTGGCAGCGGTGAATATGCCGCTGCCcctaccctacgtcctcctgc | |
| KR_SPR_wblA_5_F | GTTGGGACGTTACGGAAGGTGGGCACAGCGCAACACCCCC | 30742.72 |
| KR_SPR_wblA_5_R | GGGGGTGTTGCGCTGTGCCCACCTTCCGTAACGTCCCAACcctaccctacgtcctcctgc | |
| KR_SPR_WblA_6_F | CAGCGCAACACCCCCTTCGGGCCCAATCTTGGATGGCCCG | 30743.7 |
| KR_SPR_WblA_6_R | CGGGCCATCCAAGATTGGGCCCGAAGGGGGTGTTGCGCTGcctaccctacgtcctcctgc | |
| KR_SPR_WblA_7_F | ATCTTGGATGGCCCGAACGGACTATGCGTGCGCGTCAGAT | 30739.78 |
| KR_SPR_WblA_7_R | ATCTGACGCGCACGCATAGTCCGTTCGGGCCATCCAAGATcctaccctacgtcctcctgc | |
| KR_SPR_WblA_8_F | GCGTGCGCGTCAGATCACCCAGGGGAGTGATCGAAGGACA | 30741.74 |

| | | |
|---|---|---|
| KR_SPR_WblA_8_R | TGTCCTTCGATCACTCCCCTGGGTGATCTGACGCGCACGCcctaccctacgtcctcctgc | |
| KR_SPR_WblA_9_F | AGTGATCGAAGGACATGCGTGTGATAGCGGACATATCGCC | 30737.82 |
| KR_SPR_WblA_9_R | GGCGATATGTCCGCTATCACACGCATGTCCTTCGATCACTcctaccctacgtcctcctgc | |
| KR_SPR_WblA_10_F | AGCGGACATATCGCCTGGATGTGTCCTTCCTCGACGGGGC | 30741.74 |
| KR_SPR_WblA_10_R | GCCCCGTCGAGGAAGGACACATCCAGGCGATATGTCCGCTcctaccctacgtcctcctgc | |
| KR_SPR_WblA_11_F | CTTCCTCGACGGGGCCACTTGGGGCACCGTGTGACGCATG | 30743.7 |
| KR_SPR_WblA_11_R | CATGCGTCACACGGTGCCCCAAGTGGCCCCGTCGAGGAAGcctaccctacgtcctcctgc | |
| KR_SPR_WblA_12_F | ACCGTGTGACGCATGAGTCGGAATC | 21469.81 |
| KR_SPR_WblA_12_R | GATTCCGACTCATGCGTCACACGGTcctaccctacgtcctcctgc | |

| Label | Sequence | ds MW (da) |
|---|---|---|
| KR_SPR_vnz_04640_1_F | GCCGCGCCCGAACACCCCGCCCGGACGGACTAGCCCACTG | 30747.62 |
| KR_SPR_vnz_04640_1_R | CAGTGGGCTAGTCCGTCCGGGCGGGGTGTTCGGGCGCGGCcctaccctacgtcctcctgc | |
| KR_SPR_vnz_04640_2_F | CGGACTAGCCCACTGGGACCACCCCGAACCACCCGAAGAG | 30743.7 |
| KR_SPR_vnz_04640_2_R | CTCTTCGGGTGGTTCGGGGTGGTCCCAGTGGGCTAGTCCGcctaccctacgtcctcctgc | |
| KR_SPR_vnz_04640_3_F | GAACCACCCGAAGAGACGGTGCAGTTGACCTGAACCGCCC | 30741.74 |
| KR_SPR_vnz_04640_3_R | GGGCGGTTCAGGTCAACTGCACCGTCTCTTCGGGTGGTTCcctaccctacgtcctcctgc | |
| KR_SPR_vnz_04640_4_F | TGACCTGAACCGCCCCTGCGGGTGATGCGTCTTCCCAGGA | 30742.72 |

| | | |
|---|---|---|
| KR_SPR_vnz_04640_4_R | TCCTGGGAAGACGCATCACCCGCAGGGGCGGTTCAGGTCAcctaccctacgtcctcctgc | |
| KR_SPR_vnz_04640_5_F | TGCGTCTTCCCAGGAAAGGACGAGCTGCCGCGAAATCCCT | 30740.76 |
| KR_SPR_vnz_04640_5_R | AGGGATTTCGCGGCAGCTCGTCCTTTCCTGGGAAGACGCAcctaccctacgtcctcctgc | |
| KR_SPR_vnz_04640_6_F | TGCCGCGAAATCCCTGCGGAAGGCCGTCCATGGGGCAAGA | 30742.72 |
| KR_SPR_vnz_04640_6_R | TCTTGCCCCATGGACGGCCTTCCGCAGGGATTTCGCGGCAcctaccctacgtcctcctgc | |
| KR_SPR_vnz_04640_7_F | GTCCATGGGGCAAGACTGGGACCGGAACCCAACCGACCGA | 30742.69 |
| KR_SPR_vnz_04640_7_R | TCGGTCGGTTGGGTTCCGGTCCCAGTCTTGCCCCATGGACcctaccctacgtcctcctgc | |
| KR_SPR_vnz_04640_8_F | AACCCAACCGACCGACGGGGGCGGCGATGAGCGCGAAGAG | 30744.68 |
| KR_SPR_vnz_04640_8_R | CTCTTCGCGCTCATCGCCGCCCCCGTCGGTCGGTTGGGTTcctaccctacgtcctcctgc | |
| KR_SPR_vnz_04640_9_F | GATGAGCGCGAAGAGCCGCAGGACATCCACCACGACGCAG | 30742.7 |
| KR_SPR_vnz_04640_9_R | CTGCGTCGTGGTGGATGTCCTGCGGCTCTTCGCGCTCATCcctaccctacgtcctcctgc | |
| KR_SPR_vnz_04640_10_F | TCCACCACGACGCAGCGAAAGAACCCACCCATGTGCCAGC | 30741.8 |
| KR_SPR_vnz_04640_10_R | GCTGGCACATGGGTGGGTTCTTTCGCTGCGTCGTGGTGGAcctaccctacgtcctcctgc | |
| KR_SPR_vnz_04640_11_F | CACCCATGTGCCAGCACCAGCCTGC | 21472.77 |
| KR_SPR_vnz_04640_11_R | GCAGGCTGGTGCTGGCACATGGGTGcctaccctacgtcctcctgc | |

| Label | Sequence | ds MW (da) |
|---|---|---|
| KR_SPR_vnz_16110_1_F | TGTCGCCGCCATGTCGATTTCCTCCCGAAATCCATGTAGT | 30737.8 |

| | | |
|---|---|---|
| KR_SPR_vnz_16110_1_R | ACTACATGGATTTCGGGAGGAAATCGACATGGCGGCGACAcctaccctacgtcctcctgc | |
| KR_SPR_vnz_16110_2_F | CGAAATCCATGTAGTCAAATTTGACCACAGCAGTAGTCAA | 30733 |
| KR_SPR_vnz_16110_2_R | TTGACTACTGCTGTGGTCAAATTTGACTACATGGATTTCGcctaccctacgtcctcctgc | |
| KR_SPR_vnz_16110_3_F | CACAGCAGTAGTCAACTTTGATTACTCGCCGCCGACGGAG | 30738 |
| KR_SPR_vnz_16110_3_R | CTCCGTCGGCGGCGAGTAATCAAAGTTGACTACTGCTGTGcctaccctacgtcctcctgc | |
| KR_SPR_vnz_16110_4_F | TCGCCGCCGACGGAGCAAGTTATGCCCTCGCGCAGACTTA | 30741.72 |
| KR_SPR_vnz_16110_4_R | TAAGTCTGCGCGAGGGCATAACTTGCTCCGTCGGCGGCGAcctaccctacgtcctcctgc | |
| KR_SPR_vnz_16110_5_F | CCTCGCGCAGACTTAGTCAAACTTGATTAGCGCTGCTGAT | 30736.87 |
| KR_SPR_vnz_16110_5_R | ATCAGCAGCGCTAATCAAGTTTGACTAAGTCTGCGCGAGGcctaccctacgtcctcctgc | |
| KR_SPR_vnz_16110_6_F | ATTAGCGCTGCTGATGCGCCATGCT | 21469.78 |
| KR_SPR_vnz_16110_6_R | AGCATGGCGCATCAGCAGCGCTAATcctaccctacgtcctcctgc | |

| Label | Sequence | ds MW (da) |
|---|---|---|
| KR_SPR_vnz_16115_1_F | CGTGCTGCTTCACCGCGCCCAGGACGAGCAGCCGGATCGC | 30745.66 |
| KR_SPR_vnz_16115_1_R | GCGATCCGGCTGCTCGTCCTGGGCGCGGTGAAGCAGCACGcctaccctacgtcctcctgc | |
| KR_SPR_vnz_16115_2_F | GAGCAGCCGGATCGCCGACATCGCCGTACCTCCTAGTCAA | 30741.74 |
| KR_SPR_vnz_16115_2_R | TTGACTAGGAGGTACGGCGATGTCGGCGATCCGGCTGCTCcctaccctacgtcctcctgc | |
| KR_SPR_vnz_16115_3_F | GTACCTCCTAGTCAATTTTGACCAGGGTACGCGCGGCGGC | 30740.76 |

| | | |
|---|---|---|
| KR_SPR_vnz_16115_3_R | GCCGCCGCGCGTACCCTGGTCAAAATTGACTAGGAGGTACcctaccctacgtcctcctgc | |
| KR_SPR_vnz_16115_4_F | GGTACGCGCGGCGGCGGGGGCCGGGCGGATCAGCCCTGC | 30750.59 |
| KR_SPR_vnz_16115_4_R | GCAGGGCTGATCCCGCCCGGCCCCCGCCGCCGCGCGTACCcctaccctacgtcctcctgc | |
| KR_SPR_vnz_16115_5_F | CGGGATCAGCCCTGCCCCTGCTCCG | 21474.71 |
| KR_SPR_vnz_16115_5_R | CGGAGCAGGGGCAGGGCTGATCCCGcctaccctacgtcctcctgc | |

| Label | Sequence | ds MW (da) |
|---|---|---|
| KR_orrA_Promoter_SPR_For_1 | CCG GCC CGA AGC TGT CCG CCA TCG TTC CTC CCC CTG AAG G | 30744.68 |
| KR_orrA_Promoter_SPR_Rev_1 | CCT TCA GGG GGA GGA ACG ATG GCG GAC AGC TTC GGG CCG GCC TAC CCT ACG TCC TCC TGC | |
| KR_orrA_Promoter_SPR_For_2 | TCC TCC CCC TGA AGG CCG TGG CCT GAA TTC GTT TCC GGT C | 30741.74 |
| KR_orrA_Promoter_SPR_Rev_2 | GAC CGG AAA CGA ATT CAG GCC ACG GCC TTC AGG GGG AGG ACC TAC CCT ACG TCC TCC TGC | |
| KR_orrA_Promoter_SPR_For_3 | AAT TCG TTT CCG GTC CTC CAA CCG CTA CTC AGG GAG CAC C | 30739.78 |
| KR_orrA_Promoter_SPR_Rev_3 | GGT GCT CCC TGA GTA GCG GTT GGA GGA CCG GAA ACG AAT TCC TAC CCT ACG TCC TCC TGC | |
| KR_orrA_Promoter_SPR_For_4 | TAC TCA GGG AGC ACC GGT TGG CGT GCA CGT CAT GAT TTC A | 30738.8 |
| KR_MtrD_Promoter_SPR_Rev_4 | TGA AAT CAT GAC GTG CAC GCC AAC CGG TGC TCC CTG AGT ACC TAC CCT ACG TCC TCC TGC | |
| KR_orrA_Promoter_SPR_For_5 | CAC GTC ATG ATT TCA TGC CCA GGC GGC AAC GGC GCG ATC C | 30741.74 |
| KR_orrA_Promoter_SPR_Rev_5 | GGA TCG CGC CGT TGC CGC CTG GGC ATG AAA TCA TGA CGT GCC TAC CCT ACG TCC TCC TGC | |
| KR_orrA_Promoter_SPR_For_6 | GCA ACG GCG CGA TCC GCA GGC GCA CGA GGG TGC CCC TGG G | 30747.62 |

| | | |
|---|---|---|
| KR_orrA_Promoter_SPR_Rev_6 | CCC AGG GGC ACC CTC GTG CGC CTG CGG ATC GCG CCG TTG CCC TAC CCT ACG TCC TCC TGC | |
| KR_orrA_Promoter_SPR_For_7 | GAG GGT GCC CCT GGG GGG CGC CAG AGG CGC TCC AGG GGC A | 30748.6 |
| KR_orrA_Promoter_SPR_Rev_7 | TGC CCC TGG AGC GCC TCT GGC GCC CCC CAG GGG CAC CCT CCC TAC CCT ACG TCC TCC TGC | |
| KR_orrA_Promoter_SPR_For_8 | GGC GCT CCA GGG GCA CCG TGA GGG GCG AGG GGA GGG TGG G | 30748.64 |
| KR_orrA_Promoter_SPR_Rev_8 | CCC ACC CTC CCC TCG CCC CTC ACG GTG CCC CTG GAG CGC CCC TAC CCT ACG TCC TCC TGC | |
| KR_orrA_Promoter_SPR_For_9 | CGA GGG GAG GGT GGG CGC TGA AGC GGC CGT CAG CCG CCG A | 30747.62 |
| KR_orrA_Promoter_SPR_Rev_9 | TCG GCG GCT GAC GGC CGC TTC AGC GCC CAC CCT CCC CTC GCC TAC CCT ACG TCC TCC TGC | |
| KR_orrA_Promoter_SPR_For_10 | GCC GTC AGC CGC CGA GCG CAC CTC C | 21475.69 |
| KR_orrA_Promoter_SPR_Rev_10 | GGA GGT GCG CTC GGC GGC TGA CGG CCC TAC CCT ACG TCC TCC TGC | |

| Label | Sequence | ds MW (da) |
|---|---|---|
| wbla_7_ABO_1_F | ATCTTGGATCTCATGACTGAACTCTGCGTGCGCGTCAGAT | 30736.84 |
| wbla_7_ABO_1_R | ATCTGACGCGCACGCAGAGTTCAGTCATGAGATCCAAGATcctaccctacgtcctcctgc | |
| wbla_7_ABO_2_F | ATCTTGGAGTTAAATCCATTCAGCTGCGTGCGCGTCAGAT | 32558.05 |
| wbla_7_ABO_2_R | ATCTGACGCGCACGCAGCTGAATGGATTTAACTCCAAGATcctaccctacgtcctcctgc | |
| VNZ16110_2_ABO_1_F | CGAAATCCATGTCTCATGACTGAACTCCAGCAGTAGTCAA | 30734.89 |
| VNZ16110_2_ABO_1_R | TTGACTACTGCTGGAGTTCAGTCATGAGACATGGATTTCGcctaccctacgtcctcctgc | |
| VNZ16110_2_ABO_2_F | CGAAATCCATGGCTGACCCGGGTCAACCAGCAGTAGTCAA | 30738.8 |

| | | |
|---|---|---|
| VNZ16110_2_ABO_2_R | TTGACTACTGCTGGTTGACCCGGGTCAGCCATGGATTTCGcctaccctacgtcctcctgc | |
| vnz16115_3_ABO_1_F | GTACCTCCTCTCATGACTGAACTCGGGTACGCGCGGCGGC | 30742.72 |
| vnz16115_3_ABO_1_R | GCCGCCGCGCGTACCCGAGTTCAGTCATGAGAGGAGGTACcctaccctacgtcctcctgc | |
| vnz16115_3_ABO_2_F | GTACCTCCGCTGACCGGGGTCAACGGGTACGCGCGGCGGC | 32240.6 |
| vnz16115_3_ABO_2_R | GCCGCCGCGCGTACCCGTTGACCCCGGTCAGCGGAGGTACcctaccctacgtcctcctgc | |
| vnz04640_2_ABO_1_F | AACTCTCTCATGACTGAACTCCCCCGAACCACCCGAAGAG | 30738.8 |
| vnz04640_2_ABO_1_R | CTCTTCGGGTGGTTCGGGGGAGTTCAGTCATGAGAGAGTTcctaccctacgtcctcctgc | |
| vnz04640_2_ABO_2_F | TCAACGCTAAACAGTTTCAACCCCCGAACCACCCGAAGAG | 30737.82 |
| vnz04640_2_ABO_2_R | CTCTTCGGGTGGTTCGGGGGTTGAAACTGTTTAGCGTTGAcctaccctacgtcctcctgc | |
| vnz_04640_1_ABO_1_F | GCCGCGCCCGAACACCGAGTTCAGTCATGAGATCTCATGA | 30739.78 |
| vnz_04640_1_ABO_1_R | TCATGAGATCTCATGACTGAACTCGGTGTTCGGGCGCGGCcctaccctacgtcctcctgc | |
| vnz_04640_1_ABO_2_F | GCCGCGCCCGAACACCAATAAATTCATTCAGCTAAACAGT | 30735.86 |
| vnz_04640_1_ABO_2_R | ACTGTTTAGCTGAATGAATTTATTGGTGTTCGGGCGCGGCcctaccctacgtcctcctgc | |

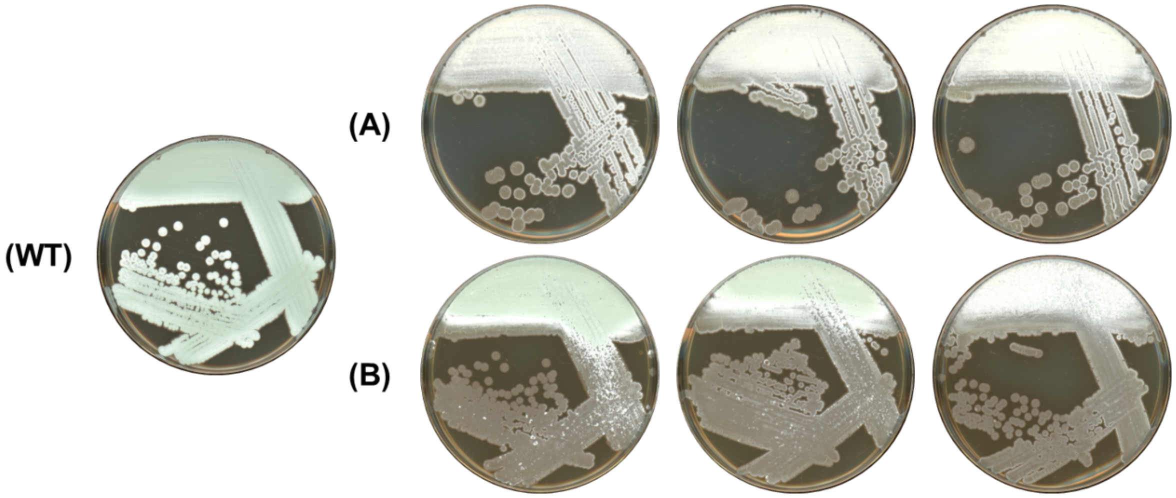


**Figure S1. The *S. venezuelae* Δ*orrA* mutant is unstable.** (A). *S. venezuelae* Δ*orrA* strains independently propagated from vegetative stocks on MYM agar. (B). *S. venezuelae* Δ*orrA* strains propagated from emergent aerial hyphae appearing after 4 days of growth on MYM. Emergent aerial hyphae are genetically unstable and revert towards a wildtype phenotype. Wild type control (WT) shown for reference. All strains were grown for three days before imaging.

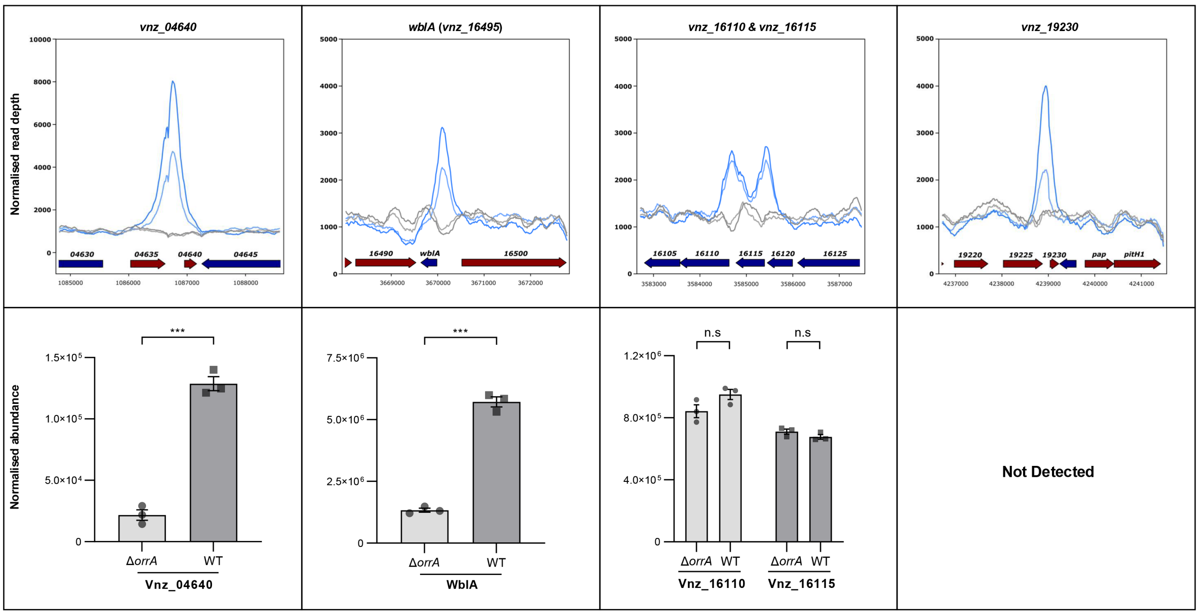


**Figure S2. OrrA ChIP-seq targets in *S. venezuelae.*** Top. Panels show the OrrA ChIP-seq peaks in blue and wild-type controls in black. Bottom. TMT proteomics data comparing abundance levels of ChIP-seq target gene products between Δ*orrA* and wild-type *S. venezuelae*. The *vnz_19320* gene product was not detected in the TMT proteomics experiment.

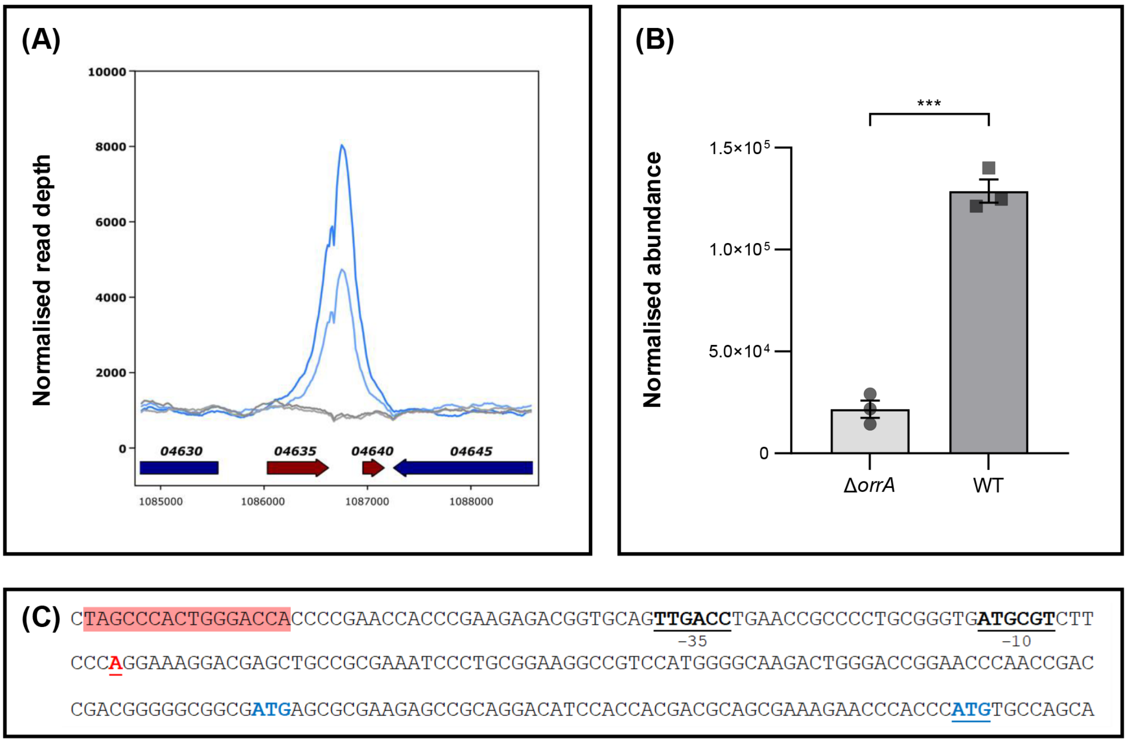


**Figure S3. OrrA directly activates the production of Vnz_04640 in *Streptomyces venezuelae*.** (A). Normalised ChIP-seq read depth, blue traces show the OrrA-Flag strain (n=2) and black traces show the wild-type control (n=2). (B). TMT proteomics data (n=3) showing the normalized peptide abundance of Vnz_04640 in the *S. venezuelae* Δ*orrA* and wild-type strains at 18 hours growth on MYM medium. Individual replicate abundances are shown. *** = BH-adjusted $p$-value < $1 \times 10^{-7}$ (background-based $t$-test). (C). The *vnz_04640* promoter with OrrA consensus site highlighted in red, the RNA polymerase -35 and -10 sites underlined in black boldface, and the transcription start site (TSS) underlined in red boldface. There are two possible Vnz_04640 translational start codons that are in frame, both are marked in blue with the annotated start codon underlined.

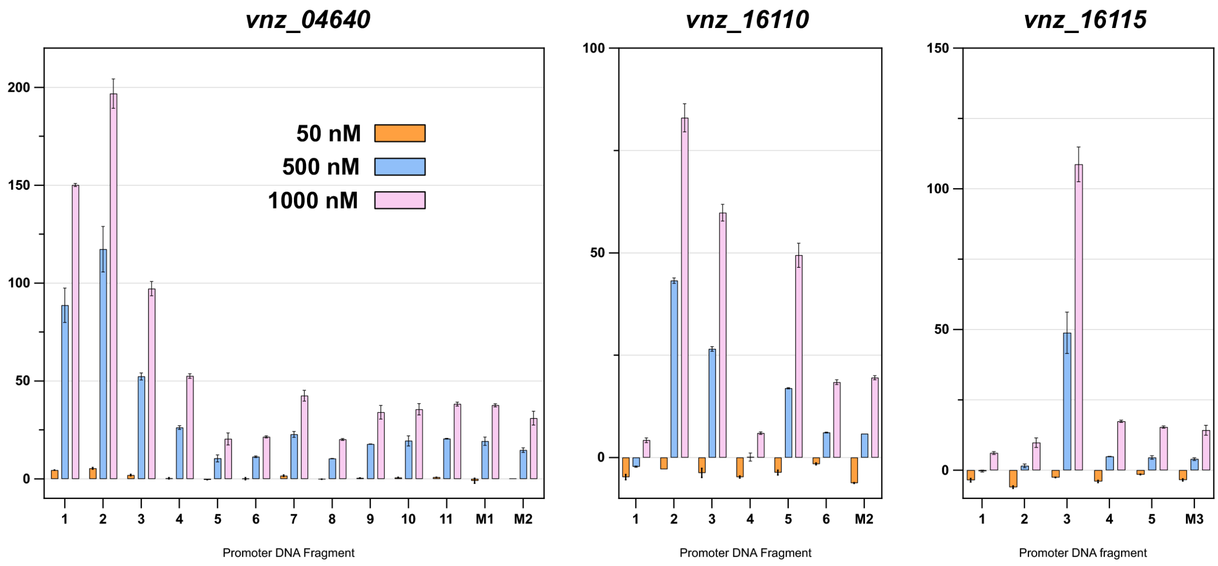


**Figure S4. OrrA binding to ChIP-seq target promoters as detected by ReDCaT SPR.** These data show binding of the purified OrrA DNA binding domain to double stranded oligonucleotide probes that tile across the target promoters. Binding is represented as $\%R_{Max}$ values calculated assuming two OrrA-DBDs binding to the imperfect palindromic MEME consensus. Error bars shown (± SEM, n = 2). The key shows concentrations of oligos used in each experiment. Oligos labelled 'M' have mutated OrrA binding sites where key residues were altered (Table S4), numbers (M1-M3) denote the probe with prominent $\%R_{Max}$ values from which they are derived. The OrrA binding sites are reported in Table 1.

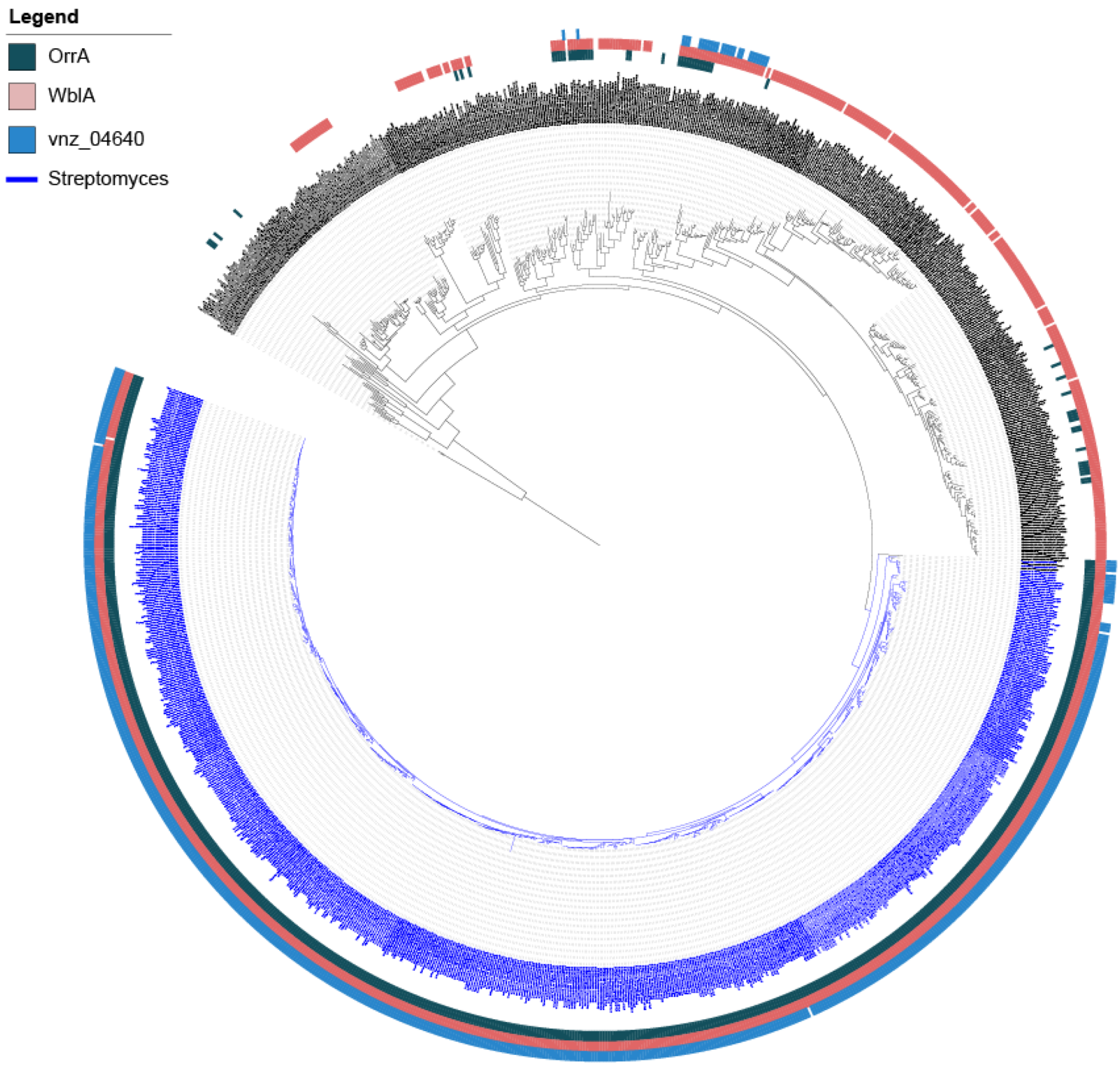


**Figure S5. Phylogenetic analysis of Actinomycetota species with presence/absence of OrrA, WblA and Vnz_04640**. Maximum likelihood phylogenetic tree of species from the phylum Actinomycetota based on MAFFT alignment of RpoB amino acid sequences. Branches and branch labels in blue indicate *Streptomyces* species. The outer rings, from inside to out, show the presence or absence of OrrA (green), WblA (pink) and Vnz_04640 (blue) homologues. The tree was constructed using FASTTREE for maximum likelihood inference and visualised in iTOL.

**Supplementary Methods (TMT Proteomics)**

Protein pellets were resuspended in 100 µl of 2.5% sodium deoxycholate (SDC; Merck) in 0.2 M EPPS-buffer (Merck), pH 8.5, and vortexed and heated to ~90°C for a total of three cycles. Protein concentration was estimated using a BCA assay and approx. 200 µg of protein per sample was reduced, alkylated, and digested with trypsin in the SDC buffer according to standard procedures. After the digest, the SDC was precipitated by adjusting to 0.2% trifluoroacetic acid (TFA), and the clear supernatant subjected to C18 SPE (Reprosil, Dr. Maisch GmbH, Germany). Peptide concentration was further estimated by running an aliquot of the digests on LCMS (methods see below). TMT labelling was performed using 6 channels (see Table 1 below) from a TMT™16plex kit (Lot WJ329096, ThermoFisher Scientific) according to the manufacturer's instructions with slight modifications; approx. 100 µg of the dried peptides were dissolved in 90 µl of 0.2 M EPPS buffer (Merck)/10% acetonitrile, and 250 µg TMT reagent dissolved in 22 µl of acetonitrile was added. After 2 h incubation, aliquots of 2 µl from each sample were combined in 400 µl 0.2% TFA, desalted, and analysed on the mass spectrometer (same method as below for TMT, but without RTS) to check labelling efficiency and estimate total sample abundances. The main sample aliquots were then quenched by adding 8 µl of 5% hydroxylamine and then combined correspondingly to roughly level abundances. The combined samples were desalted using a C18 Sep-Pak cartridge (200 mg, Waters), redissolved in 500 µl of 25 mM $NH_4HCO_3$ and fractionated by high pH reversed phase HPLC. Using an ACQUITY Arc Bio System (Waters), the samples were loaded to an XBridge® 5 µm BEH C18 130 Å column (250 x 4.6 mm, Waters). Fractionation was performed with the following gradient of solvents A (water), B (acetonitrile), and C (25 mM $NH_4HCO_3$ in water) at a flow rate of 1 ml min$^{-1}$: solvent C was kept at 10% throughout the gradient; solvent B: 0-5 min: 5%, 5-10 min: 5-10%, 10-80 min: 10-45%, 80-90 min: 45-80%, followed by 5 min at 80% B and re-equilibration to 5% for 24 min. Fractions of 1 ml were collected and concatenated by combining fractions of similar peptide concentration to produce 19 final fractions for MS analysis. Aliquots were analysed by nanoLC-MS/MS on an Orbitrap Eclipse™ Tribrid™ mass spectrometer equipped with a FAIMS Pro Duo interface coupled to an UltiMate® 3000 RSLCnano LC system (Thermo Fisher Scientific, Hemel Hempstead, UK). The samples were loaded onto a trap cartridge (Pepmap 100, C18, 5um, 0.3x5mm, Thermo) with 0.1% TFA at 15 µl min$^{-1}$ for 3 min. The trap column was then switched in-line with the analytical column. A nanoEase M/Z column (HSS C18 T3, 1.8 µm, 100 Å, 250 mm x 0.75 µm, Waters) was used for separation using the following gradient of solvents A (water, 0.1% formic acid) and B (80% acetonitrile, 0.1% formic acid) at 40°C at a flow rate of 0.2 µl min$^{-1}$ : 0-3 min 3% B (parallel to trapping); 3-10 min linear increase B to 8 % (curve 4); 10-108 min linear increase B to 50%; 108-113 min linear increase B to 99 %; kept at 99% B for 3 min and re-equilibration to 3% B. Data were acquired on the mass spectrometer with the following parameters in positive ion mode: MS1/OT: resolution 120K, profile mode, mass range *m/z* 400-1600, AGC target 4e$^5$, max inject time 50 ms, FAIMS device set to three compensation voltages (-35V, -50V, -65V) for 1 s each; MS2/IT: for each CV, data dependent analysis with the following parameters: 1 s cycle time Rapid mode, centroid mode, quadrupole isolation window 0.7 Da, charge states 2-5, threshold 1.9e$^4$, CID CE = 30, AGC target 1e4, max. inject time 50 ms, dynamic exclusion 1 count for 15 s mass tolerance of 7 ppm; MS3 synchronous precursor selection (SPS): 10 SPS precursors, isolation window 0.7 Da, HCD fragmentation with CE=50, Orbitrap Turbo TMT and TMTpro resolution 30k, AGC target 1e5, max inject time 100 ms, Real Time Search (RTS): protein database *Streptomyces venezuelae* (7420 entries, from *Streptomyces venezuelae* strain NRRL B-65442 chromosome - Nucleotide - NCBI (nih.gov)), enzyme trypsin, 1 missed cleavage, oxidation (M) as variable, carbamidomethyl (C) and TMTpro as fixed modifications, precursor tolerance 10 ppm, Xcorr = 1.4, dCn = 0.1.

The mass spectrometry raw data were processed and quantified in Proteome Discoverer 3.2 (Thermo Fisher Scientific); all mentioned tools of the following workflow are nodes of the proprietary Proteome Discoverer (PD) software. The *S. venezuelae* fasta database (as above for RTS) was imported into PD adding a reversed sequence database for decoy searches; a database for common contaminants (maxquant.org, 245 entries) was also included. The database search was performed using the incorporated search engines CHIMERYS (MSAID, Munich, Germany) and Comet (Eng, Jahan and Hoopmann, 2013). The processing workflow included recalibration (RC) of MS1 spectra, reporter ion quantification by most confident centroid (20 ppm) and a search on the imported *S. venezuelae* database. The Top N Peak Filter was applied with 20 peaks per 100 Da. For CHIMERYS, the inferys_4.7.0._fragmentation prediction model was used with fragment tolerance of 0.3 Da, enzyme trypsin with 1 missed cleavage, variable modification oxidation (M), fixed modifications carbamidomethyl (C) and TMTpro on N-terminus and K. For Comet the version 2019.01 rev.0 parameter file was used with default settings except precursor tolerance set to 6 ppm and trypsin missed cleavages set to 1. Modifications were the same as for CHIMERYS.

The consensus workflow included the following parameters: intensity-based abundance, normalisation on total peptide abundances, protein abundance-based ratio calculation, only unique peptides (protein groups) for quantification, TMT channel correction values applied (according to the TMT kit lots mentioned above), co-isolation/SPS matches/CHIMERYS Coefficient thresholds 50%/65%, 0.8, missing values imputation by low abundance resampling, hypothesis testing by t-test (background based), adjusted p-value calculation by Benjamini-Hochberg method. The results were exported into a Microsoft Excel table including data for normalised and un-normalised abundances, ratios for the specified conditions, the corresponding p-values and adjusted p-values, number of unique peptides, q-values, PEP-values,  identification scores from both search engines; FDR confidence filtered for high confidence (strict FDR 0.01) only.

### REFERENCES


Cobb, R.E., Wang, Y. and Zhao, H. (2015) 'High-Efficiency Multiplex Genome Editing of *Streptomyces* Species Using an Engineered CRISPR/Cas System', *ACS Synthetic Biology*, 4(6), pp. 723–728. Available at: https://doi.org/10.1021/sb500351f.

Eng, J.K., Jahan, T.A. and Hoopmann, M.R. (2013) 'Comet: An open-source MS / MS sequence database search tool', *Proteomics*, 13(1), pp. 22–24. Available at: https://doi.org/10.1002/pmic.201200439.

Gomez-Escribano, J.P. *et al.* (2021) '*Streptomyces venezuelae* NRRL B-65442: genome sequence of a model strain used to study morphological differentiation in filamentous actinobacteria', *Journal of Industrial Microbiology and Biotechnology*, 48(9–10), p. kuab035. Available at: https://doi.org/10.1093/jimb/kuab035.

Hong, H.-J. *et al.* (2005) 'The Role of the Novel Fem Protein VanK in Vancomycin Resistance in Streptomyces coelicolor', *Journal of Biological Chemistry*, 280(13), pp. 13055–13061. Available at: https://doi.org/10.1074/jbc.M413801200.

Jordan, M.L. and Schlimpert, S. (2025) 'Microbe Profile: Streptomyces venezuelae – a model species to study morphology and differentiation in filamentous bacteria. *Microbiology*, 171(3). Available at: https://doi.org/10.1099/mic.0.001541.

Schlimpert, S. *et al.* (2017) 'Two dynamin-like proteins stabilize FtsZ rings during Streptomyces sporulation', *Proceedings of the National Academy of Sciences*, 114(30), pp. E6176–E6183.

Schlimpert, S. and Elliot, M.A. (2023) 'The Best of Both Worlds—Streptomyces coelicolor and Streptomyces venezuelae as Model Species for Studying Antibiotic Production and Bacterial Multicellular Development', *Journal of Bacteriology*, pp. e00153-23. Available at: https://doi.org/10.1128/jb.00153-23.
